# Multi-modal adaptor-clathrin contacts drive coated vesicle assembly

**DOI:** 10.1101/2021.05.22.445240

**Authors:** Sarah M. Smith, Gabrielle Larocque, Katherine M. Wood, Kyle L. Morris, Alan M. Roseman, Richard B. Sessions, Stephen J. Royle, Corinne J. Smith

**Affiliations:** School of Life Sciences, University of Warwick, Gibbet Hill Road, Coventry CV4 7AL, UK; Centre for Mechanochemical Cell Biology, Warwick Medical School, University of Warwick, Gibbet Hill Road, Coventry CV4 7AL, UK; Division of Molecular and Cellular Function, School of Biological Sciences, Faculty of Biology, Medicine and Health, University of Manchester, Manchester Academic Health Science Centre, Manchester M13 9PT, UK; School of Biochemistry, University of Bristol, Medical Sciences Building, University Walk, Bristol BS8 1TD, UK; Cellular Signalling and Cytoskeletal Function Laboratory, The Francis Crick Institute, London NW1 1AT, UK; Diamond Light Source Ltd, Harwell Science & Innovation Campus, Didcot, UK

**Keywords:** clathrin, endocytosis, membrane traffic, cryo-electron microscopy

## Abstract

Clathrin-coated pits are formed by the recognition of membrane and cargo by the heterotetrameric AP2 complex and the subsequent recruitment of clathrin triskelia. A potential role for AP2 in coated-pit assembly beyond initial clathrin recruitment has not been explored. Clathrin binds the β2 subunit of AP2, and several binding sites on β2 and on the clathrin heavy chain have been identified, but our structural knowledge of these interactions is incomplete and their functional importance during endocytosis is unclear. Here, we analysed the cryo-EM structure of clathrin cages assembled in the presence of β2 hinge and appendage (β2HA) domains. We find that the β2-appendage binds in at least two positions in the cage, demonstrating that multi-modal binding is a fundamental property of clathrin-AP2 interactions. In one position, β2-appendage cross-links two adjacent terminal domains from different triskelia below the vertex. Functional analysis of β2HA-clathrin interactions reveals that endocytosis requires two clathrin interaction sites: a clathrin-box motif on the hinge and the “sandwich site” on the appendage, with the appendage “platform site” having less importance. From these studies and the work of others, we propose that β2-appendage binding to more than one clathrin triskelion is a key feature of the system and likely explains why clathrin assembly is driven by AP2.

## Introduction

Clathrin-mediated endocytosis (CME) is the major route of entry for receptors and their ligands into cells (Mettlen et al., 2018). A clathrin-coated pit is formed at the plasma membrane that selects cargo for uptake into the cell via a clathrin-coated vesicle. Clathrin cannot recognize membrane or cargo itself and so an adaptor protein binds the membrane, selects the cargo, and associates with clathrin leading to pit formation (Figure 1A). Several adaptor proteins have clathrin binding sites and colocalize with clathrin structures in cells but the assembly polypeptide-2 (AP2) complex (α, β2, µ2, and σ2 subunits) is thought to primarily initiate clathrin recruitment. The recruitment of clathrin by the β2 subunit is an essential step in CME. AP2 and clathrin arrive jointly at the membrane in a ratio of two AP2 complexes per triskelion (Cocucci et al., 2012). As the pit matures, the ratio decreases as clathrin polymerizes (Bucher et al., 2018). It is assumed that this polymerization – which is an innate property of clathrin triskelia – completes vesicle formation. However, AP2 is named after its ability to promote clathrin cage assembly *in vitro* (Pearse and Robinson, 1984; Zaremba and Keen, 1983), and a fragment of the β2 subunit of AP2, containing the hinge and appendage domains (β2HA), has been shown to promote the polymerization of clathrin (Gallusser and Kirchhausen, 1993; Owen et al., 2000; Shih et al., 1995). How these *in vitro* observations relate to endocytosis in cells is unclear. One intriguing but often overlooked idea is that AP2, via β2HA, serves a dual role in CME: initially recruiting clathrin to the plasma membrane and then driving coated vesicle assembly. There are two clathrin-binding locations on β2HA (Figure 1B). The first is a linear peptide motif within the hinge region (Lundmark and Carlsson, 2002; Owen et al., 2000), LLNLD, called the clathrin box motif (CBM). The second clathrin-binding location is within the β2-appendage domain, however its precise nature is debated (Chen and Schmid, 2020). The appendage domain has two sites that interact distinctly with different binding partners (Edeling et al., 2006; Owen et al., 2000; Schmid et al., 2006). The first, termed the sandwich (or side) domain, which surrounds Tyr 815, binds AP180, amphiphysin and eps15. A second site, termed the platform (or top) domain, surrounds residues Y888 and W841. This binds the adaptor proteins epsin, β-arrestin and autosomal recessive hypercholesterolemia (ARH) protein and functions independently from the sandwich domain. The roles of these sites in clathrin binding remain to be clarified. *In vitro* pull-down experiments highlight the potential importance of both Y888 and Y815 for clathrin binding but reports differ on their relative contribution (Edeling et al., 2006; Owen et al., 2000; Schmid et al., 2006).

**Figure 1.**
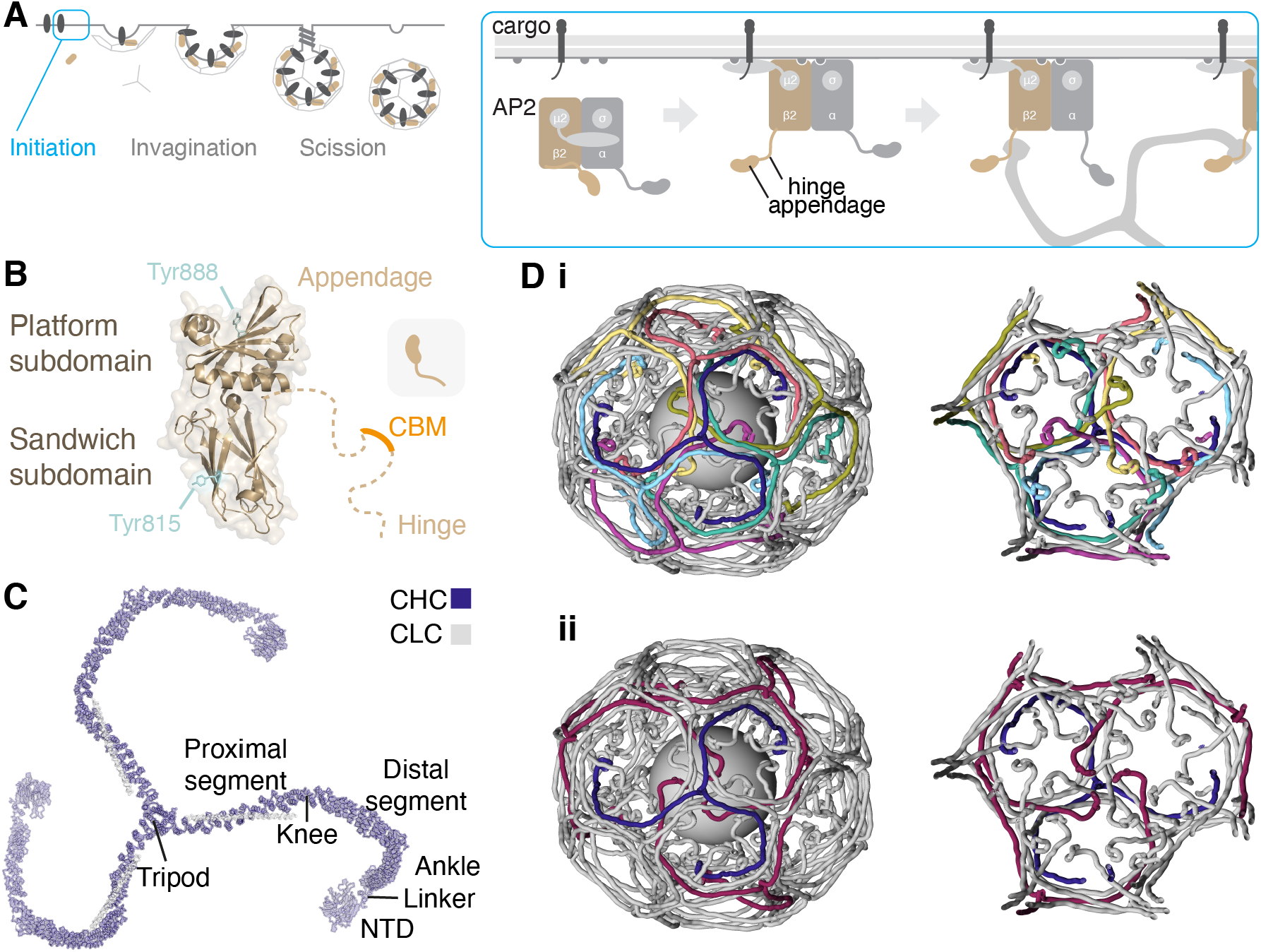
Structural view of clathrin assembly during endocytosis. (**A**) Schematic diagram of clathrin-mediated endocytosis. The AP2 complex opens when it engages cargo and PI(4,5)P2, the β2 hinge and appendage (β2HA) become available for clathrin binding, initiating pit formation. (**B**) Structure of β2HA (PDB code: 2G302G30). The appendage is divided into platform and sandwich subdomain, each with a tyrosine residue previously identified to be important for clathrin binding. The unstructured hinge region contains a clathrin-box motif (CBM, LLNLD) which binds the N-terminal domain (NTD) of clathrin heavy chain. (**C**) Structure of a clathrin triskelion (PDB code: 3IYV). Three clathrin heavy chains (CHC) each with an associated light chain (CLC) are trimerized at their C-termini forming a tripod. Each leg is divided into proximal and distal segments, an ankle region and NTD. (**D**) Clathrin assemblies. i) An indigo triskelion is shown engaged with six other triskelia in a hexagonal barrel, coating a vesicle. Each edge is made from four leg segments for four different triskelia: two antiparallel proximal regions on the outer surface and two antiparallel distal regions below. ii) The tripod of this triskelion is at a vertex, and below that, three NTDs are arranged, contributed by triskelia (purple) whose tripods are two or three edges away. Right panels show the view from the vesicle toward the vertex. The positions of triskelia were mapped by downsampling the carbon backbones in 3IYV by 5 residues and smoothing their position in 3D space using a 25 residue window in IgorPDB. CLCs have been removed for clarity.

Our structural understanding of how clathrin engages with AP2 is incomplete. The N-terminal domain (NTD, Figure 1C) of clathrin heavy chain is a seven-bladed β-propeller with four adaptor protein binding sites (Willox and Royle, 2012). Atomic structures have revealed that CBMs bind promiscuously to these sites, with the AP2 CBM binding to the “CBM site” between blades 1 and 2 and also to the “arrestin site” between blades 4 and 5 (Muenzner et al., 2017). The location where β2-appendage binds clathrin is uncertain. Knuehl *et al*. used biochemical approaches and yeast-2-hybrid studies to identify residues C682 and G710 on the heavy chain ankle region as a potential location for β2-appendage (Knuehl et al., 2006). Another potential location is where transforming acidic coiled-coil 3 (TACC3) binds clathrin (residues 457-507, (Burgess et al., 2018; Hood et al., 2013)). However a full picture of how the β2HA interacts with assembled clathrin, central to the mechanism of clathrin recruitment, remains elusive.

Recently, two structural studies have visualized contradictory modes of binding for the β2-appendage in clathrin assemblies. Using cryo-electron tomography, Kovtun *et al*. investigated the structure of assembled clathrin and a form of AP2 lacking the alpha appendage and hinge region on lipid membranes containing cargo peptides and PI(4,5)P2 (Kovtun et al., 2020). They observed density beneath the clathrin vertex enclosed by one terminal domain and the ankle regions of two triskelion legs. In contrast, Paraan *et al*. isolated native coated vesicles from bovine brain and obtained a structure using single particle analysis. They observed density consistent with the β2-appendage, however it was in a different location, between two adjacent terminal domains (Paraan et al., 2020). In order to address the paradox, we have analyzed the structure of purified clathrin bound to the β2HA using single particle cryo-EM approaches. We find that the β2-appendage binds in at least two positions on clathrin, within the same sample, demonstrating that multi-modal binding is a fundamental property of clathrin-AP2 interactions and reconciling the differing observations in the literature. Our functional analysis of β2HA-clathrin interactions reveals that endocytosis requires hinge and appendage interaction sites, with the Tyr 815 sandwich site being more important for productive vesicle formation than the Tyr 888 platform site. In consolidating all available structural and functional information, we find that β2-appendage binding to more than one clathrin triskelion is a key feature of the system and likely explains how clathrin assembly is driven by AP2.

## Results

### The appendage of β2 is critical for coated vesicle formation

We previously developed a strategy to trigger clathrin-coated vesicle formation in cells, termed “hot-wired endocytosis” (Wood et al., 2017). It works by inducibly attaching a clathrin-binding protein (clathrin “hook”) to a plasma membrane “anchor” using an FKBP-rapamycin-FRB dimerization system; and this is sufficient to trigger endocytosis (Figure 2A). Using the hinge and appendage of the β2 subunit of the AP2 complex (FKBP-β2HA-GFP) as a clathrin hook allows us to examine endocytosis that is driven by the interaction of β2HA and clathrin, that is, independent of other clathrin-adaptor interactions. Hot-wired endocytosis can be detected in live cells by visualizing the formation of intracellular bright green puncta that also contain an antibody to the extracellular portion of the anchor. Using FKBP-β2HA-GFP as a clathrin hook, the formation of numerous puncta was observed, while a control construct (FKBP-GFP) elicited no response (Figure 2B,C).

**Figure 2.**
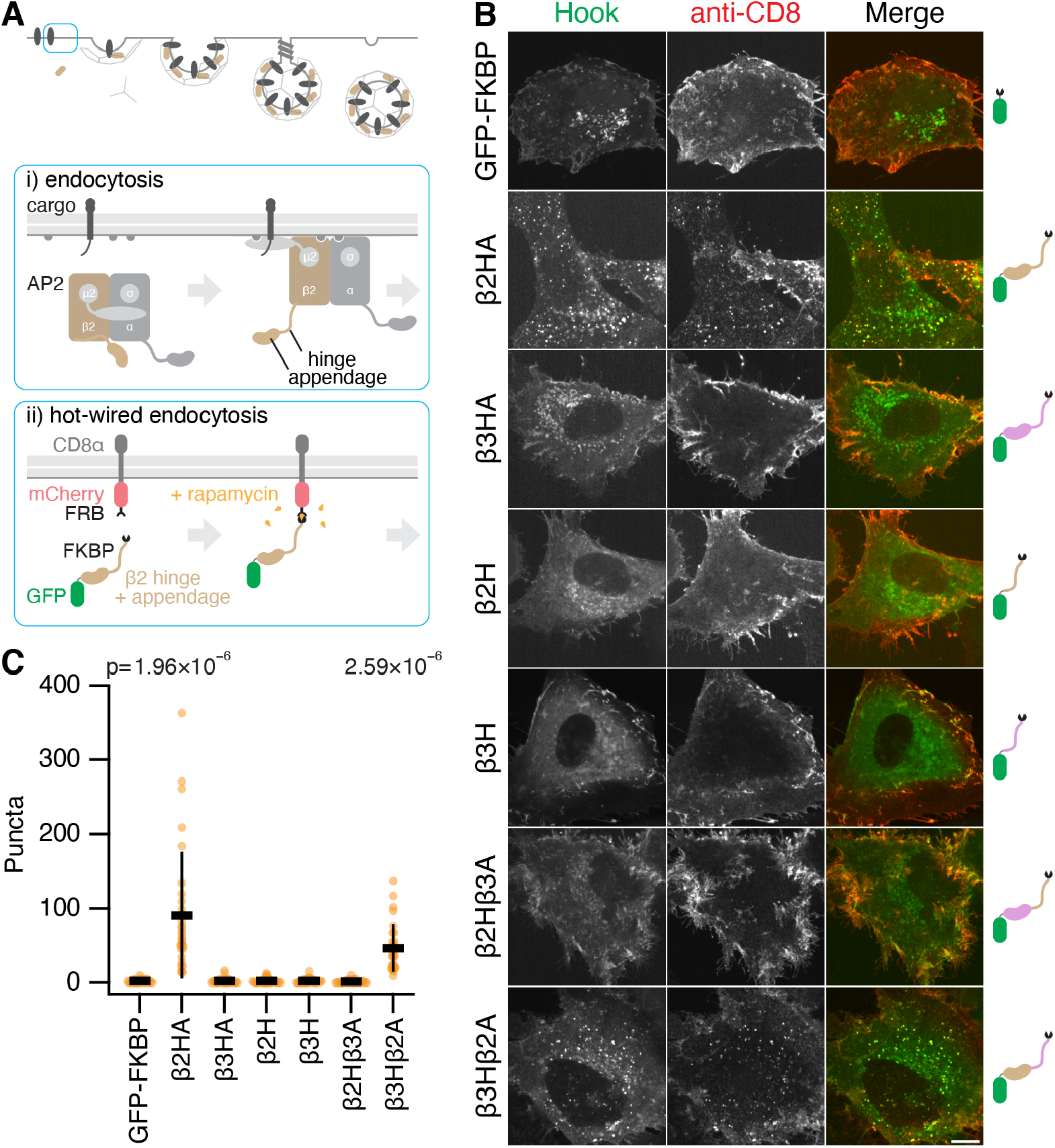
The β2 appendage is crucial for hot-wired clathrin mediated endocytosis. (**A**) Schematic diagram of hot-wired endocytosis. Under normal conditions (i) the AP2 complex engages with cargo and PI(4,5)P2 and the β2 hinge and appendage become available for clathrin engagement; in hot-wiring (ii) a clathrin hook, e.g. β2 hinge and appendage (FKBP-β2HA-GFP) is attached to a plasma membrane anchor CD8-mCherry-FRB inducibly by rapamycin application. (**B**) Representative confocal micrographs of cells expressing the plasma membrane anchor (CD8-mCherry-FRB) and the indicated hooks (green). Cells were incubated with anti-CD8-Alexa647 (red) and treated with rapamycin (200 nM, 15 min). Vesicles coinciding in both green and red channels (yellow in merge) were quantified in B. Scale bar, 10 µm. (**C**) Scatter dot plot shows the number of intracellular GFP-positive vesicles that contained anti-CD8 Alexa647 per cell, bars indicate mean ± SD. Number of experiments = 3. P-values from Dunnett’s post-hoc test that were less than 0.1 are shown above.

An analogous construct from the AP3 complex, FKBP-β3HA-GFP, with the hinge and appendage of β3, was not competent for hot-wiring (Figure 2B,C). This is a surprising result for two reasons: first, the clathrin box motif in the hinge of β3 binds clathrin *in vitro* (Dell’Angelica et al., 1998), and second, we had assumed that the role of the clathrin hook in the hot-wiring system was solely to recruit clathrin initially, with downstream polymerization being driven by clathrin alone.

To investigate this result in more detail, we tested whether the hinges of β2 or β3 were competent for hot-wiring. Despite the presence of a clathrin box motif in both hinges, with the appendage domains removed neither FKBP-β2H-GFP nor FKBP-β3H-GFP was able to induce endocytosis (Figure 2B,C). Next, we transplanted the appendage of β3 onto the β2 hinge, and the appendage of β2 onto the β3 hinge. We observed hot-wiring with FKBP-β3Hβ2A-GFP but not with FKBP-β2Hβ3A-GFP (Figure 2B,C). Thus the β2 appendage was able to drive endocytosis with a β3 hinge but the β2 hinge alone or in the presence of the β3 appendage could not. These results indicate firstly that the β2 appendage is critical for endocytosis and that the β3 appendage cannot substitute for this activity. Secondly, hooks containing a clathrin-box motif are not sufficient for vesicle formation. This suggested to us that the β2 appendage is active in clathrin polymerization.

### Structure of clathrin-β2HA minicoat cages

If the β2 appendage contributes to clathrin polymerization, the nature of its interaction with assembled clathrin is of particular interest. In order to investigate this we analyzed cryo-electron micrographs of clathrin assembled in the presence of β2HA (Figure S1). Saturation of β2HA binding sites on clathrin was achieved using a 60-fold molar excess of β2HA (Figure S1A,B). Of the 57 528 particles analyzed, 29 % of the total particle data set (16 641 particles) was occupied by the minicoat class of cages (Figure S1F,G). Subsequent extensive supervised and unsupervised 3D classifications identified the particles most stably identified with the minicoat cage architecture (Figures S2 and S3). These 13 983 minicoat particles were refined to a gold-standard resolution of 9.1 Å (Figure S4).

In order to locate β2HA within the map density we compared the β2HA-clathrin map to a map of clathrin cages assembled in the absence of β2HA. While a difference map did reveal density in a location just above the terminal domains, it was not well-defined (Figure 3A,B). We therefore conducted a voxel-by-voxel comparison between the two maps to locate statistically significant differences (Young et al., 2013). This method allows the location of differences to be determined with confidence but does not define the shape of difference density. This enabled us to evaluate the entire minicoat particle dataset globally for potential β2HA binding locations. The results of this analysis confirmed a significant difference just above the terminal domains (Figure 3C). We also noted significant differences in some other areas, away from β2HA, that may be related to triskelion leg movements or other conformational changes upon β2HA binding.

**Figure 3.**
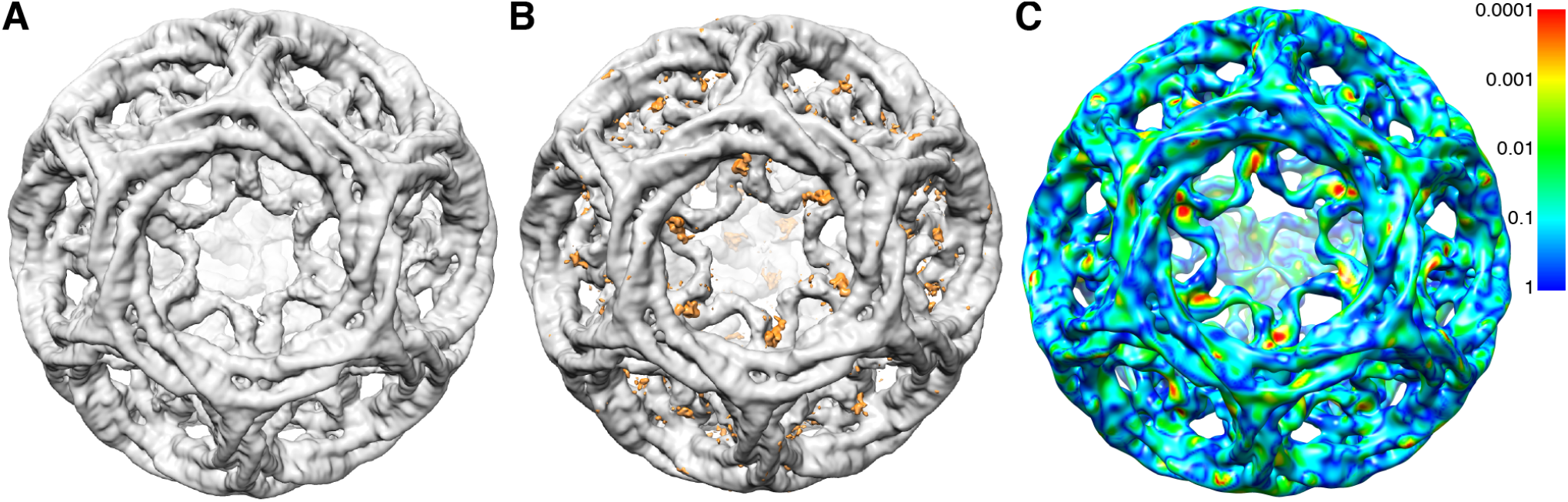
Global difference analysis of clathrin-β2HA compared to clathrin alone. (**A**) Unsharpened cryo-EM map of clathrin-β2HA minicoat cage architecture at 9.1 Å resolution. (**B**) Difference map of clathrin-β2HA minicoat and clathrin-only minicoat. Differences in density are shown in orange. (**C**) Clathrin-β2HA and clathrin-only minicoat maps. Statistically significant differences are shown on a rainbow color scheme (see inserted panel) with red, orange, yellow and green being the areas of significant difference. The light blue and dark blue areas indicate regions where the significance is below our threshold, or there is no significant difference between the two maps. The regions with the most significant difference density at *p* < 0.0005 (in red) were interpreted as the binding site of β2HA. Other regions show significant differences due to conformation changes related to binding. The contour level of all maps is 3.0 times sigma above the mean of the map. All images were created in UCSF Chimera (Pettersen et al., 2004).

### Finding β2HA in clathrin-β2HA minicoats

Our global difference analysis suggested that the β2HA was indeed bound to the cages but not well-resolved due to heterogeneity. The flexibility of clathrin terminal domains, which results in weaker density (Fotin et al., 2004; Morris et al., 2019), likely contributes to the heterogeneity. We used signal subtraction to reduce the dominance of the strong features of the outer clathrin cage in order to classify the weaker terminal domain signal more precisely (Bai et al., 2015) (Figure S5). 13 983 particles of the inner region of the minicoat cage were classified into 20 classes, with occupancy ranging from 1.4 % to 12.2 %, reflecting the heterogeneity of this cage region. Particles belonging to each class were refined individually to a higher resolution (Figure S6). The outputs of the individual refinements (each at contour level σ3) varied in the quality and completeness of the terminal domain density. However in two classes, 15 and 18, distinct density consistent with bound β2HA was observed.

In the case of class 15, these densities were in a different location to that shown by our global analysis, on alternate terminal domains within a polyhedral face (Figure S7). Comparison of equivalent positions in a minicoat cage without adaptor bound demonstrate that the densities present at the terminal domains were a consequence of β2HA binding (Figure S7). Looking at adjacent polyhedral faces, for a given hub region where 3 terminal domains (from separate triskelia) converge, two terminal domains are engaged in an interaction with a single β2-appendage leaving one terminal domain unoccupied (Figure S7). Interestingly, β2HA density was not present at any of the 4 hubs in the minicoat cage where 3 pentagonal faces join. This class was refined further using localized reconstruction (described below).

For class 18, density could be seen on every terminal domain in all the hexagonal faces of the minicoat volume, but was less well-resolved (Figure 4A,B). In contrast to class 15 these densities lay parallel to the terminal domain beta-propeller and did not contact neighboring terminal domains. We used localized reconstruction (Ilca et al., 2015; Morris et al., 2019) to improve the resolution of the hexagonal faces from this class of minicoat particles. Rigid-body fitting of the clathrin terminal domain atomic structure revealed surplus density on either side of the beta propeller structure (Figure 4C). The location of this density is consistent with our earlier global difference analysis. The surplus density at either side of the terminal domain was large enough to accommodate the atomic structure of the β2-appendage (Figure 4D) but could not support an unambiguous fit of this structure.

**Figure 4.**
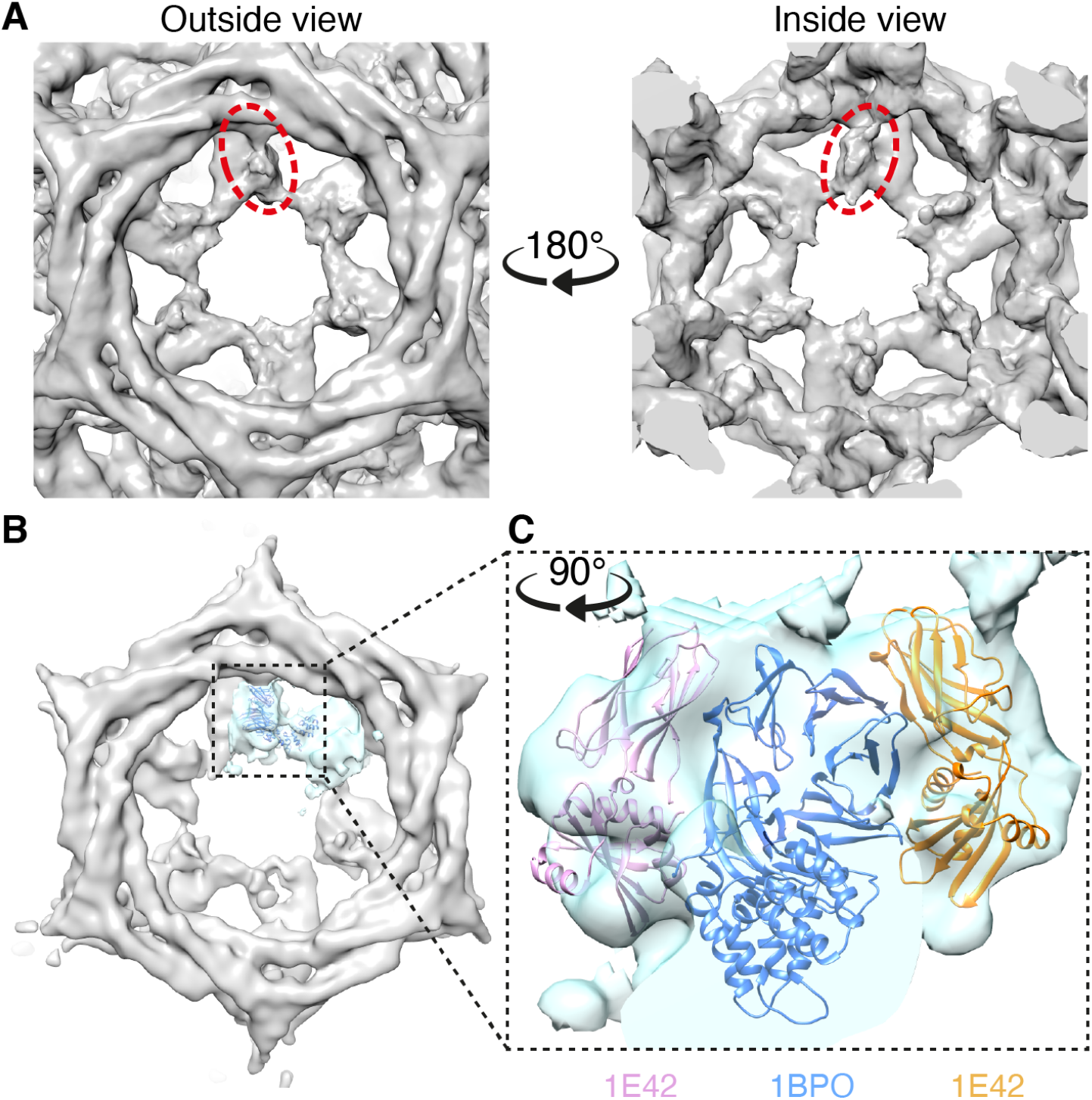
Unattributed density in hexagonal faces of clathrin minicoats. (**A**) Close-up view of the unattributed density on each terminal domain in a given hexagonal face of the clathrin-β2HA minicoat cage that was resolved from particles belonging to class 18. Views from outside and inside a given hexagonal face are shown (left and right, respectively). Example densities are highlighted in dashed red ellipses. (**B**) Using localized reconstruction (Ilca et al., 2015), all hexagonal faces from the minicoat cage shown in panel A were extracted and averaged resulting in a 19 Å map. A single terminal domain and connecting linker and ankle region, is highlighted in cyan, with the atomic model of clathrin terminal domain β-propeller structure (PDB 1BPO) rigid body fitted into the density (shown in dark blue). (**C**) Rigid-body fitting of the clathrin terminal domain atomic structure (shown in dark blue) revealed surplus density on either side of the β-propeller structure, which was large enough to accommodate the atomic structure of the β2-adaptin appendage (PDB 1E42, shown in pink and orange) but not sufficiently defined to support an unambiguous fit to the density. All images were created, and rigid-body fitting was conducted, in UCSF Chimera (Pettersen et al., 2004).

### Resolving β2HA in the minicoat hub substructure

Having established through our analysis of whole cages that β2HA has at least two different binding locations on assembled clathrin, we next improved the resolution of the most defined density for β2HA by making use of the local symmetry present within the cages. We extracted and refined the hub regions from each vertex of the minicoat cage particles belonging to Class 15 (Figure S6), using localized reconstruction within Relion (Ilca et al., 2015). Using this approach we refined a total of 26 624 minicoat hub regions to a global resolution of 9.6 Å (Figure S8). This resulted in a considerable improvement in resolution when compared to the whole-cage particles of Class 15 which refined to 19.8 Å (Figure S6). Hubs surrounded by three pentagonal faces, which did not show additional density, were excluded from this refinement. A difference map and statistical comparison confirmed the presence of density due to β2HA (Figure 5). We also found that separating the hubs according to whether the β2HA density linked terminal domains emerging from two pentagonal faces or from one pentagonal face and one hexagonal face resulted in improved map definition (Figure S7C,D). These two classes were refined separately to global resolutions of *∼* 10 Å (10.5 Å for P-P hubs and 10.1 Å for H-P hubs, Figure S8), consistent with the reduced number of particles in each subset. Despite this slightly lower resolution, the β2HA density in these maps was more clearly defined than in previous maps, with an intensity equal to the adjoining terminal domains (at contour level σ3)), and a 1:2 β2HA:terminal domain binding ratio for both hub volumes (Figure S8A-C).

**Figure 5.**
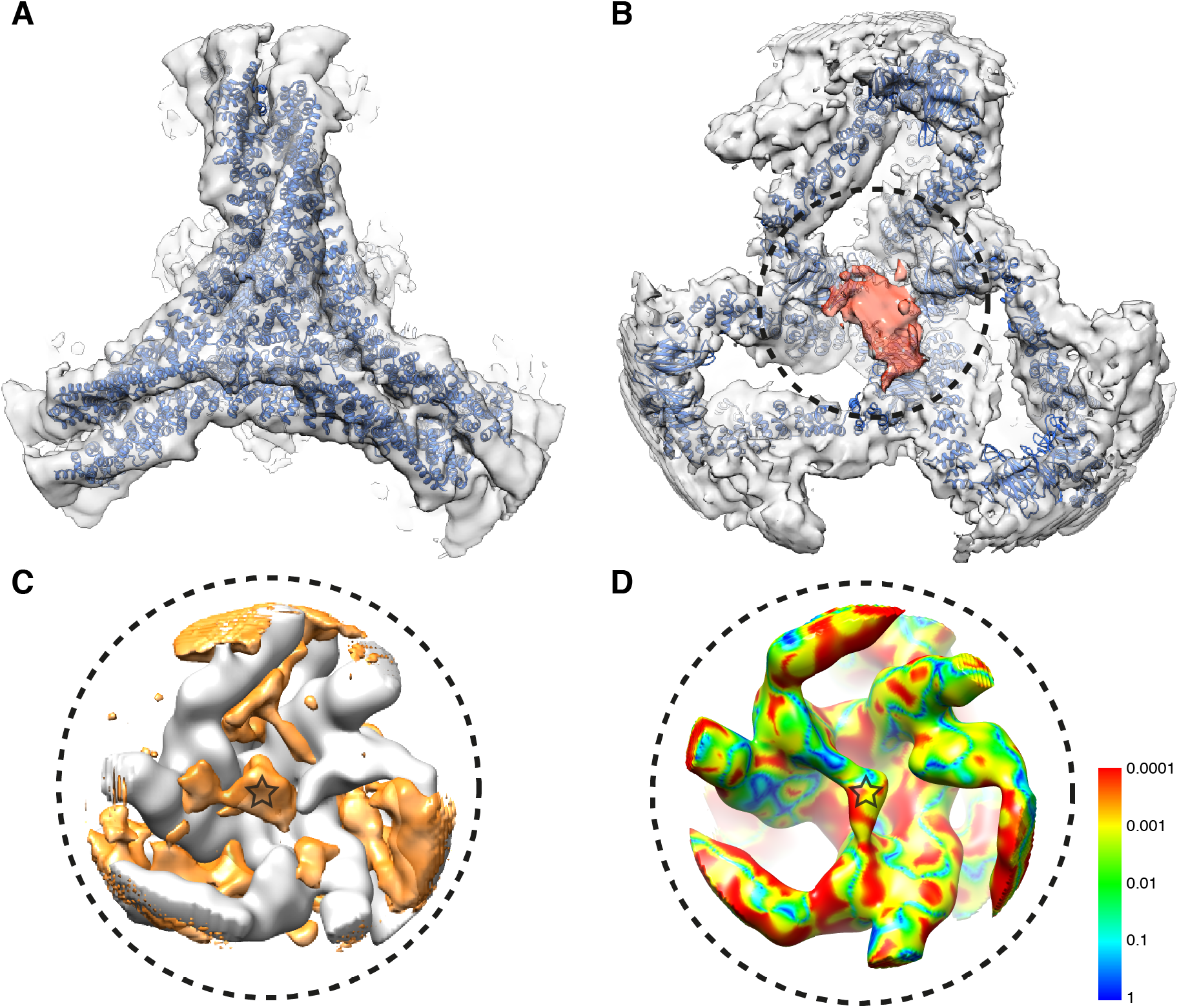
Identification of β2HA in clathrin minicoats. (**A**) 10.5 Å resolution cryo-EM map of clathrin-β2HA minicoat hub region particles belonging to class 15. The hub region atomic model (PDB 6SCT) was flexibly fitted (Joseph et al., 2016; Topf et al., 2008) into the cryo-EM map (blue). (**B**) Underside view of the cryo-EM map and fitted clathrin model shown in panel A. Density attributed to β2HA is colored red. (**C**) Difference map of the clathrin-β2HA minicoat and clathrin-only minicoat hub maps. Differences in density are shown in orange. The orange density located at the junction of the three terminal domains (marked with a star) is consistent with the location of β2HA shown in panel B. (**D**) Pixel by pixel comparison between clathrin-β2HA and clathrin-only minicoat hub maps. Statistically significant differences are shown on a rainbow color scheme (see inserted panel) with red, orange, yellow and green being the areas of significant difference. Red indicates the regions with significant differences at the highest level, *p* < 0.0005. Differences reflect the binding of β2HA and induced movements of the legs. The light blue and dark blue areas indicate regions where the significance is below our threshold, or there is no significant difference between the two maps. The density attributed to β2HA is marked with a star.

### Defining β2-appendage interactions with clathrin terminal domains

The improved definition of the β2HA density in the P-P hubs allowed us to carry out rigid-body fitting of the atomic structures of β2-appendage (PDB 1E42)(Owen et al., 2000) and clathrin terminal domain (PDB 1BPO)(ter Haar et al., 1998). The optimal orientation of the β2-appendage was found by selecting the fit with the greatest occupation of density (Figure 6A,C). A molecular model of the clathrin heavy chain formed from the model of Morris *et al*. (6SCT) and the terminal domain X-ray structure of ter Haar *et al*. (1BPO) was fitted into the P-P hub map using a combination of manual fitting and FlexEM (Morris et al., 2019; ter Haar et al., 1998; Joseph et al., 2016; Topf et al., 2008). Based on this fit, the alignment of the β2-appendage to potential binding sites on the terminal domain was then optimised according to predicted intermolecular interaction energies calculated using the programme BUDE to determine a plausible binding interface (Figure 6D,E). The resulting model has been deposited as 7OM8.pdb.

**Figure 6.**
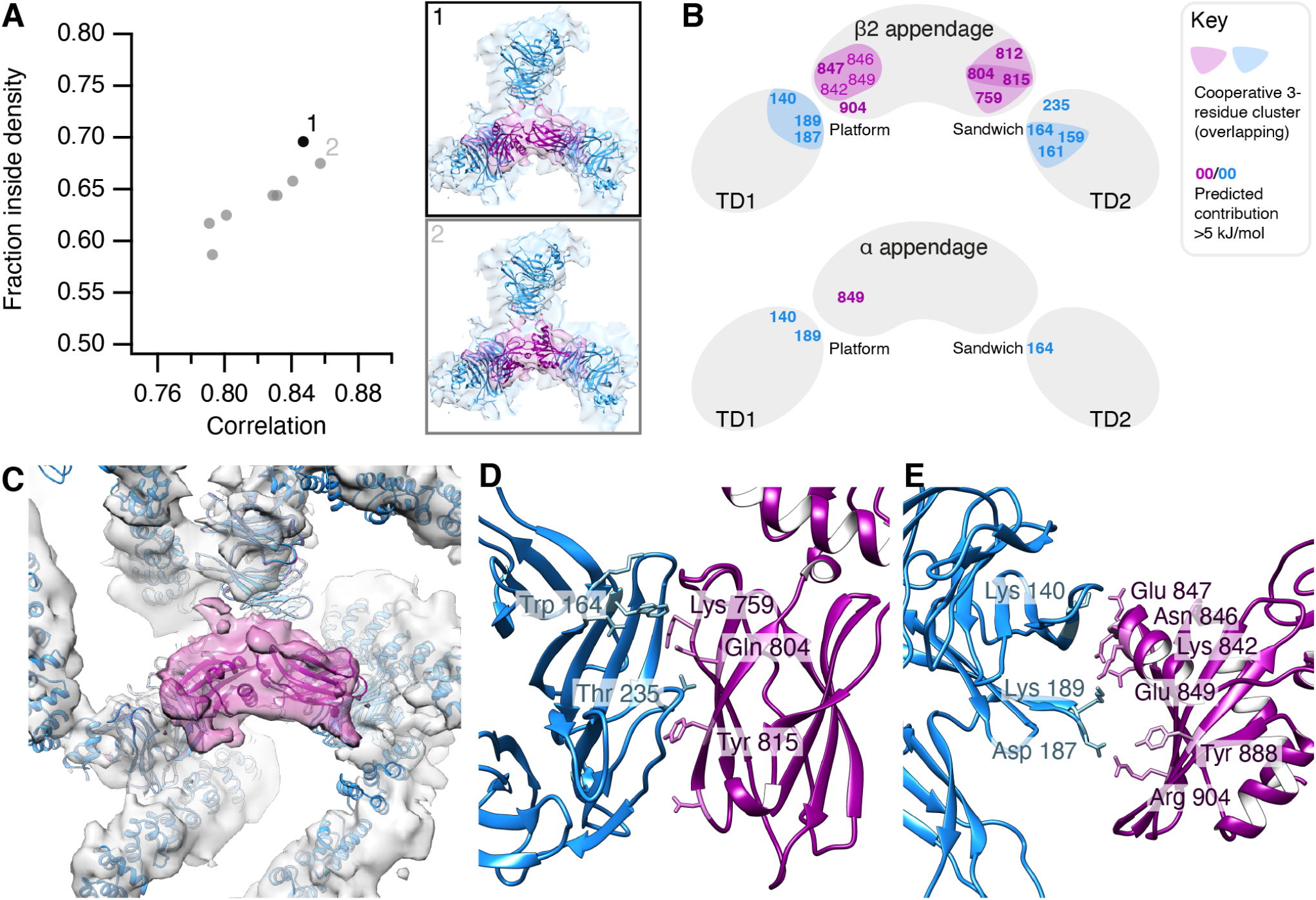
Orientation of β2-appendage with respect to clathrin terminal domains. (**A**) Plot of fraction of atomic structure inside the cryo-EM density (Y-axis) versus correlation of fit (X-axis). The fit with the largest fraction inside density is shown in black. Although it had the second highest correlation value, this orientation yielded the highest hit-rate in the rigid body fitting, accounting for 34 % of possible fits calculated. Two possible orientations (panels 1 and 2) of the β2-appendage were determined through rigid body fitting of the atomic structure of Owen *et al*., PDB 1E42 into the P-P minicoat hub volume from class 15. (**B**) Diagrammatic summary of the analysis of binding interfaces using BudeAlaScan (BAlaS). Full results are given in Table 1. (**C**) The selected best fit of the β2-appendage (purple) is shown in the context of the surrounding clathrin legs (Model in blue, density in grey). (**D**) Predicted interface between terminal domain β-propeller (blue) and β2-appendage sandwich or side domain (purple) surrounding Tyr 815. (**E**) Predicted interface between terminal domain β-propeller (blue) and β2-appendage platform or top domain (purple) surrounding Tyr 888.

We then conducted a systematic analysis of the potential contribution residues at the interface made to binding energy using the programme BudeAlaScan (BAlaS) (Ibarra et al., 2019; Wood et al., 2020) which performs computational alanine scanning (Table 1). In addition to looking at single residues we examined the effect of multiple weaker interactions to define residue clusters that, through a cooperative effect, may prove important for binding. As a control we performed a similar analysis with the α-appendage domain which does not bind clathrin. These results predict that residue Tyr 815 on the β2-appendage makes the largest contribution (14 kJ mol^−1^) with Asp 812, Gln 804 and Lys 759 contributing at or above a 5 kJ mol^−1^ threshold at that interface. There were also contributions from Glu 847 and Arg 904 at the 888 platform domain interface with a second terminal domain. Tyr 888 itself, implicated in adaptor interactions with the β2-appendage, does not form contacts with the terminal domain in our model. In the control experiments with the α-appendage, only Glu 849 showed a contribution *>*5 kJ mol^−1^ (7.2 kJ mol^−1^).

**Table 1.**
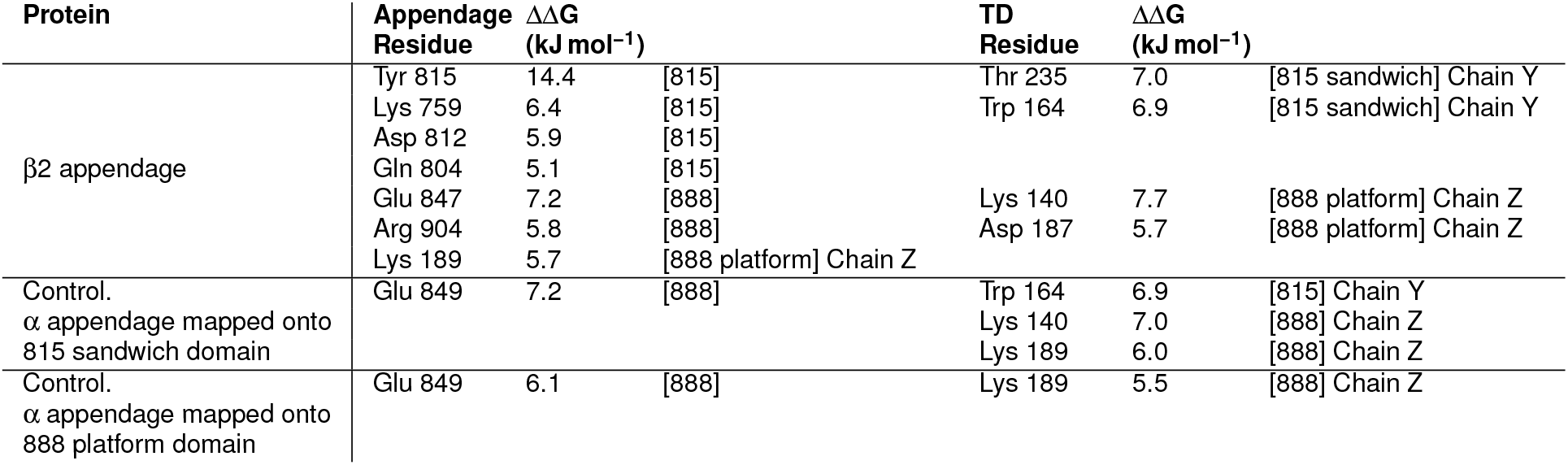
Analysis of binding interfaces using BudeAlaScan (BAlaS): alanine scanning.

A similar analysis looking at the terminal domain interactions showed only two residues contributing more than 5 kJ mol^−1^ to the 815 sandwich interface; Thr 235 and Trp 164, while three terminal domain residues contributed more than 5 kJ mol^−1^to the 888 platform domain interface; Lys 140, Asp 187 and Lys 189. In the alpha appendage controls, three residues contributed more than 5 kJ mol^−1^; Lys 140, Trp 164 and Lys 189. In all cases there were no contributions comparable to that of Tyr 815. This suggested that individual residue interactions are less important for terminal domain binding, so we investigated potential cooperativity from groups of weaker-binding residues. We performed a constellation analysis using BAlaS for residues with an interaction energy greater than 3 kJ mol^−1^. This showed that cooperative clusters formed at the interfaces between the terminal domains and both the 815 sandwich and 888 platform domains (Table 2 and Figure 6). At the 815 sandwich domain these complementary clusters involved β2 appendage residues Lys 759, Gln 804, Asp 812 and Tyr 815 and terminal domain residues Asp 159, Lys 161 and Trp 164. At the 888 platform domain interface the complementary clusters consisted of β2-appendage residues Lys 842, Asn 846, Glu 847 and Glu 849 and terminal domain residues Lys 140, Asp 187 and Lys 189. For the alpha appendage control there were no cooperative clusters at the 888 platform domain but at the 815 sandwich domain some pairs of residues showed high cooperativity (Table 2).

**Table 2.** Analysis of binding interfaces using BAlaS: constellation analysis for ΔΔG = 3 kJ mol^−1^.

| Protein | Constellation | Constellation $\Delta\Delta G$ (kJ mol<sup>-1</sup>) | Summed Individual $\Delta\Delta G$s (kJ mol<sup>-1</sup>) | Cooperativity (kJ mol<sup>-1</sup>) |
| --- | --- | --- | --- | --- |
| $\beta 2$ appendage ($\beta 2$-TD) | B759_B804_B815 | 9.7 | 25.8 | -16.2 |
|  | B804_B812_B815 | 14.1 | 25.3 | -11.2 |
|  | B842_B846_B847 | 3.8 | 15.1 | -11.3 |
|  | B842_B846_B849 | -2.4 | 12.3 | -14.7 |
|  | B842_B847_B849 | 1.5 | 15.2 | -13.6 |
|  | B846_B847_B849 | 2.3 | 16.1 | -13.8 |
| TD ($\beta 2$-TD) | Y159_Y161_Y164 | -1.4 | 14.7 | -16.1 |
|  | Z140_Z187_Z189 | 3.7 | 19 | -15.3 |
| $\alpha$ ($\alpha$ mapped onto 815 sandwich domain) | A761_A763 | -15.1 | 7.7 | -22.8 |
| TD ($\alpha$ mapped onto 815 sandwich domain) | Y161_Y164 | -13.8 | 10 | -23.8 |
|  | Z140_Z189 | -8.9 | 13 | -21.9 |
| TD ($\alpha$ mapped onto 888 platform domain) | n.a. | n.a. | n.a. | n.a. |

In all, this analysis of our proposed model suggests that Tyr 815 plays a key role in β2-appendage clathrin binding, supported by residues Asp 812, Gln 804 and Lys 759. On the terminal domain, residues Thr 235 and Trp 164, supported by Asp 159 and Lys 161 contribute to the interface (Figure 6B, D, E). It also suggests the potential for cooperative clusters of weaker binding interactions to support a binding interface between the 888 platform domain and the terminal domain.

### Role of β2 residues 815 and 888 in functional clathrin assembly

Previous work had identified Tyr 815 and Tyr 888 (shown in Figure 1B) as being important for β2HA-clathrin interactions (Edeling et al., 2006; Schmid et al., 2006). Our model and *in silico* alanine scanning analysis had identified the importance of both the platform and sandwich sites on β2-appendage in this interaction, so we returned to the hot-wired endocytosis system to address their relative functional importance. We had found that the hinge and appendage of β2 or β1 but not β3, were competent for hot-wiring (Figure 2B,C) (Wood et al., 2017). Consistent with these results, Tyr 815 and Tyr 888 are conserved in β2 and β1 but missing in β3 (Figure S9).

Deletion of the clathrin-box motif (ΔCBM) or mutation of Tyr 815 to alanine (Y815A) impaired the ability of FKBP-β2HA-GFP to generate endocytic vesicles (Figure 7). Mutation of Tyr 888 to valine (Y888V), a mutation previously reported to reduce clathrin binding (Schmid et al., 2006) had no measurable effect on hot-wiring (Figure 7). Moreover, the Y888V mutation had no additional effect when combined with ΔCBM, whereas the combination of ΔCBM and Y815A completely ablated hot-wiring (Figure 7B).

**Figure 7.**
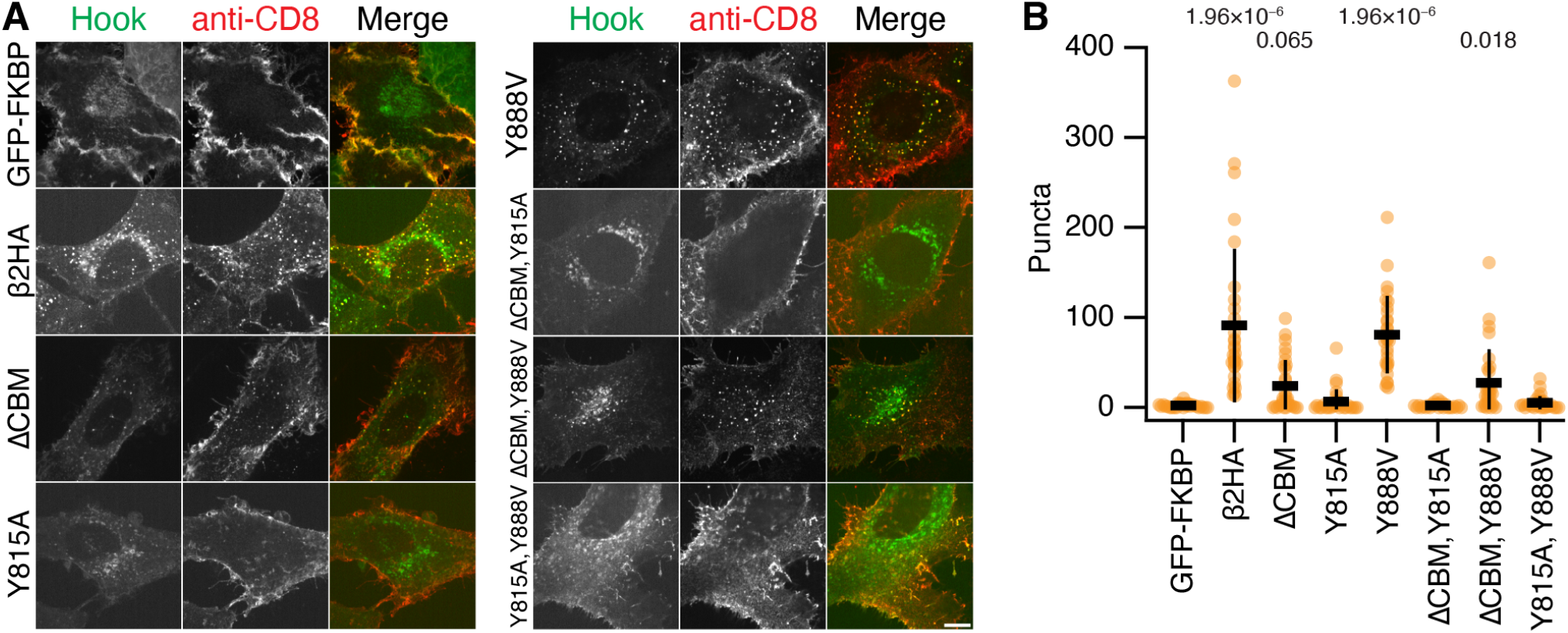
Functional test of importance of Tyr 815 and Tyr 888 to CCV formation. (**A**) Representative confocal micrographs of cells expressing the plasma membrane anchor (CD8-mCherry-FRB) and the indicated hooks (green). Cells were incubated with anti-CD8-Alexa647 (red) and treated with rapamycin (200 nM, 15 min). Vesicles coinciding in both green and red channels (yellow in merge) were quantified in B. Scale bar, 10 µm. (**B**) Scatter dot plot shows the number of intracellular GFP-positive vesicles that contained anti-CD8 Alexa647 per cell, bars indicate mean ± SD. P-values from Dunnett’s post-hoc test that were less than 0.1 are shown above. Note that these results are from the same experimental series as Figure 1, and that the negative and positive control data (GFP-FKBP and FKBP-β2HA-GFP) are as in Figure 1.

These results suggest that functional clathrin-β2HA interactions depend on the clathrin-box motif in the hinge and the sandwich site of the β2 appendage (centered on Tyr 815) while the role of the platform site of the β2 appendage (centered on Tyr 888) is undetectable in this assay.

## Discussion

In this paper we describe two positions in the clathrin cage where β2-appendage binds in the assembled state. These occur within the same sample, demonstrating that multi-modal binding is a fundamental property of clathrin-AP2 interactions. Our functional analysis demonstrated that endocytosis depends on two interactions between β2HA and clathrin. Together these observations provide an explanation for how AP2 drives coated vesicle formation in cells.

Our observation that β2HA can bind in multiple positions on clathrin within the same sample casts a new perspective on the apparently contradictory observations of Kovtun *et al*. and Paraan *et al*., suggesting they instead form part of a wider spectrum of possible β-appendage binding modes (Kovtun et al., 2020; Paraan et al., 2020). We have summarized these multi-modal clathrin-β2HA interactions in Figure 8. The first location (Class 15) is between two of the three terminal domains that sit directly beneath the vertex. The second location (Class 18) that we identified is above the terminal domain and maps closely to the location identified by Kovtun *et al*., where the appendage faces CHC repeat 2 from one triskelion and CHC repeat 1 of another. The third location, described by Paraan *et al*. is analogous to our Class 15 in that two terminal domains are linked, but they are adjacent domains within a hexagonal face rather than being beneath a cage vertex (Figure 8). A common feature of all three modes is the potential for β2HA to crosslink clathrin triskelia.

**Figure 8.**
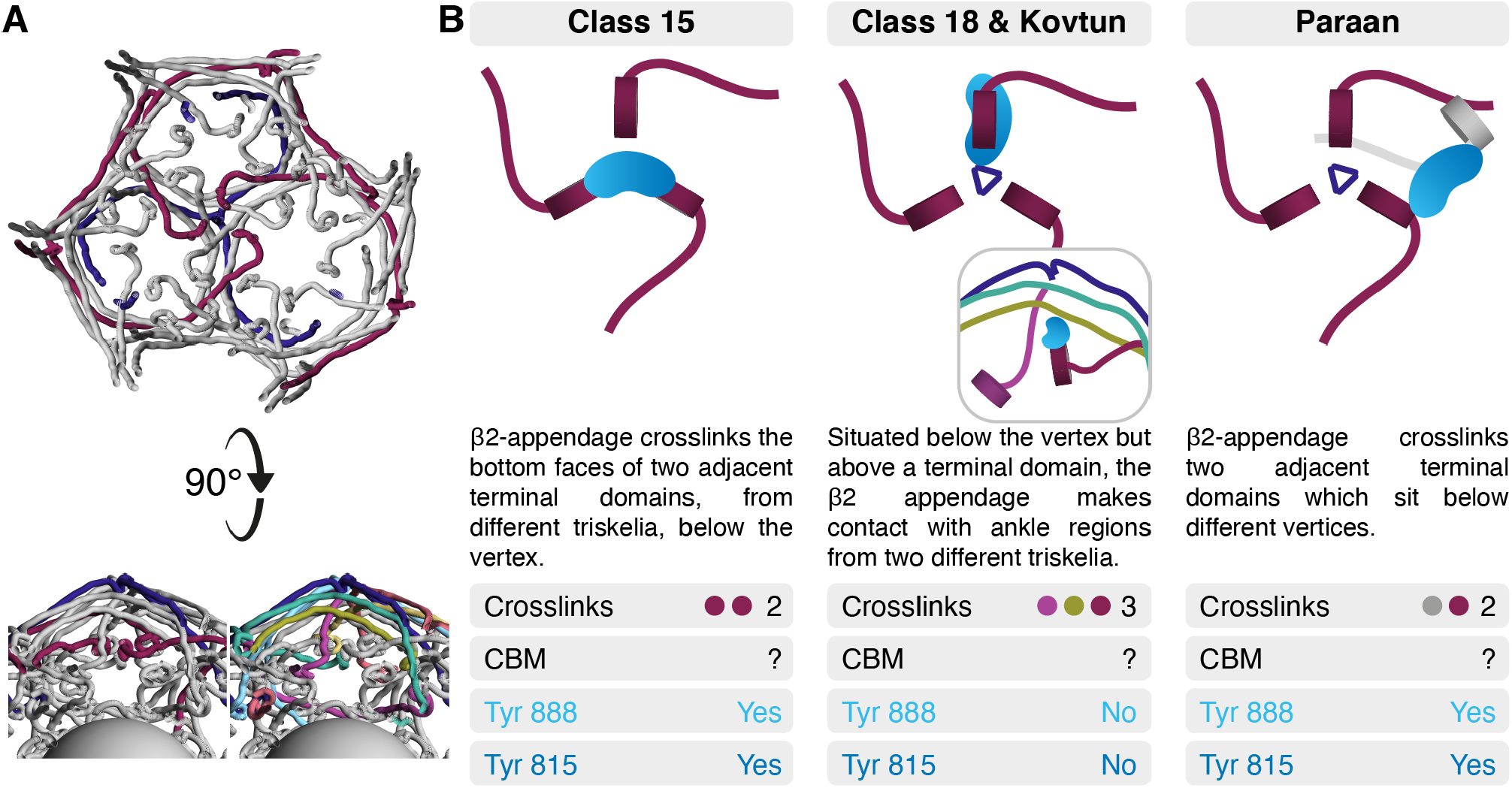
Summary of multi-modal clathrin-β2HA interactions. (**A**) For orientation, an indigo triskelion and the three NTDs situated below its vertex are shown, contributed by maroon colored triskelia (viewed from the vesicle, toward the vertex). A “side view” with the same coloring and with alternative coloring (also used in panel B) where the six triskelia that interact with the indigo triskelion are depicted (below). (**B**) Three modes of binding reported in this study and in two recent studies (Kovtun et al., 2020; Paraan et al., 2020). Common and contrasting features of each binding mode are shown. The inset in Class 18 & Kovtun shows the side view from A. Each panel indicates the number of cross links, whether CBM density was observed and whether Tyr 888 and Tyr815 are in an orientation likely to bind to the terminal domain.

In the first and third modes, the sandwich site and platform site of the β2-appendage are positioned to make interactions with two distinct terminal domains. Whereas in the second mode, according to the best fit of the β2-appendage reported by Kovtun *et al*., neither site faces clathrin; although we note that the second highest scoring fit would place the platform site in apposition to CHC repeat 2 (Kovtun et al., 2020). This region contains Cys 682 and Gly 710, previously identified as important for binding β2- and GGA-appendages (Knuehl et al., 2006).

The location of the β2-hinge is unknown in all three modes of binding. The clathrin box motif in the hinge has been shown to interact as an extended peptide slotted between blades 1 and 2 or between blades 4 and 5 of the beta-propeller at clathrin heavy chain’s N-terminal domain (Muenzner et al., 2017; ter Haar et al., 2000; Zhuo et al., 2015). At the current resolution, density relating to a peptide in such an extended conformation would be very hard to distinguish, and the promiscuous nature of clathrin box motif binding reduces the observable density further. However, we know that the interaction of this motif is essential for coated vesicle formation possibly because it is involved in initial clathrin recruitment. Assuming that it maintains contact with the terminal domain as the coat forms, all three modes of binding allow for β2HA to make contact with clathrin heavy chains from up to three distinct triskelia.

Our hot wiring results indicate that for endocytosis to proceed, the two most important sites on β2HA are the sandwich site and the clathrin box motif in the hinge, with no detectable contribution from the platform site. Previous biochemical experiments investigated the importance of Tyr 815 in the sandwich site and Tyr 888 in the platform site in the interaction of β2HA with clathrin (Edeling et al., 2006; Owen et al., 2000; Schmid et al., 2006). Owen *et al*. and Schmid *et al*. showed a significant effect on clathrin binding to the β-appendage and hinge when Tyr 888 was altered to Val (Owen et al., 2000; Schmid et al., 2006). In experiments where Tyr 815 was altered to Ala, Edeling *et al*. showed that the effect on clathrin binding was most apparent when the hinge region was either absent, or the clathrin-binding motif within the hinge was deleted or mutated (Edeling et al., 2006). While these experiments used brain extract for binding, where indirect interactions via other proteins known to bind both clathrin and β2HA remain a possibility, they nonetheless support our conclusion that the sandwich site, in partnership with the CBM domain within the β-adaptin hinge, is required for clathrin polymerization in functional coat formation. Our *in silico* analysis indicates that, in one mode, the platform site of β2HA does make contact with the terminal domain. We interpret our hot-wiring results to mean that this contact occurs only after binding of the hinge and the sandwich site. Such a cooperative role would allow interaction with other adaptor and accessory proteins such as epsin, β-arrestin and ARH at the platform domain (Schmid et al., 2006), creating flexibility to recruit the additional cargo associated with these adaptors to the growing vesicle.

All available data thus suggest that the β2 subunit of AP2 has a dual function in first recruiting clathrin to the membrane and then driving coated vesicle formation by promoting clathrin polymerization. The alternative model, where β2 only recruits clathrin and clathrin self-assembles in the absence of contribution of the adaptors is highly unlikely. First, a single clathrin box motif, which is sufficient to bind clathrin in vitro is not sufficient to induce coated vesicle formation in the hot-wiring assay. Second, under the alternative model, the requirement for the appendage in addition to the hinge would mean that both interactions must occur on one triskelion exclusively. The multiple modes of binding described from cryo-EM data all feature triskelia crosslinking, and therefore make this model improbable.

The multi-modal nature of β2HA-clathrin interactions that cross-link triskelia raise the question of whether other adaptors have the same property. Alternative adaptors such as epsin, β-arrestin and AP180 have multiple motifs that interact with clathrin or with AP2 (Smith et al., 2017; Traub, 2009). This suggests that they may also be able to contribute to initial recruitment and to cross-linking during assembly. In the case of epsin, the linear unstructured domain alone is capable of forming coated pits *in vitro* (Dannhauser and Ungewickell, 2012) and forming vesicles in cells using hot-wired endocytosis (Wood et al., 2017). The interpretation of these experiments was simply that epsin recruited clathrin and then clathrin self-assembled into a cage. It is possible that this region of epsin may also cross-link triskelia and thereby contribute to coat formation. If this is the case, it suggests a mechanism whereby adaptor proteins included in a growing coat can enhance clathrin polymerization and thereby stabilize, or even accelerate, coated vesicle formation. Since epsin, β-arrestin and AP180 also bring cargo to the growing vesicle, this has implications for understanding how particular cargos may, through their associated adaptor, increase the likelihood of completion of a growing vesicle and consequently drive forward their own internalization.

## Methods

### Molecular biology

The hinge and appendage of the human β2 subunit of the AP2 complex (designated β2HA), corresponding to residues 616-951 of the long isoform, was used for all experiments. Numbering of residues in the appendage is after the structure of β2-appendage which uses the numbering of the shorter isoform, ending at residue 937 (Owen et al., 2000).

CD8-mCherry-FRB, FKBP-β2HA-GFP, FKBP-β3HA-GFP, FKBP-β2HA(Y815A)-GFP, FKBP-β2HA(ΔCBM)-GFP and FKBP-β2HA(ΔCBM,Y815A)-GFP were described previously (Wood et al., 2017). The Y888V mutation was added by site-directed mutagenesis to FKBP-β2HA-GFP, FKBP-β2HA(Y815A)-GFP and FKBP-β2HA(ΔCBM)-GFP. FKBP-β2H-GFP was made by substituting β2-hinge (616-704) in place of β2HA in FKBP-β2HA-GFP using BamHI and AgeI. Similarly, FKBP-β3H-GFP was made by substituting β3-hinge (702-859) in FKBP-β3HA-GFP using PspOMI and AgeI. FKBP-β2Hβ3A-GFP was made by inserting the β3-appendage (860-1094) into FKBP-β2A-GFP via SalI and NotI. FKBP-β3Hβ2A-GFP was made by inserting the β2-appendage (705-951) into FKBP-β3 hinge-GFP via SalI and NotI. Plasmid to express GST-β2HA in bacteria was available from previous work (Hood et al., 2013).

### Cell culture and light microscopy

HeLa cells (Health Protection Agency/European Collection of Authenticated Cell Cultures, #93021013) were kept in DMEM supplemented with 10 % FBS and 100 U ml^−1^ penicillin/streptomycin at 37 °C and 5 % CO_2_. DNA transfection was performed with Genejuice (Merck Millipore) using the manufacturer’s protocol. HeLa cells were transfected with CD8-mCherry-FRB and one of the hooks (GFP-FKBP, FKBP-β2HA-GFP, FKBP-β3HA-GFP, FKBP-β2H-GFP, FKBP-β3H-GFP, FKBP-β2Hβ3A-GFP, FKBP-β3Hβ2A-GFP, FKBP-β2HA(ΔCBM)-GFP, FKBP-β2 HA(Y815A)-GFP, FKBP-β2HA(Y888V)-GFP, FKBP-β2HA(ΔCBM,Y815A)-GFP, FKBP-β2HA(ΔCBM,Y888V)-GFP or FKBP-β2HA(Y815A,Y888V)-GFP). After 24 h, the cells were put on coverslips. The next day, surface CD8 was labeled with 10 µg ml^−1^ AlexaFluor647-anti-CD8 antibody (Bio-Rad, MCA1226A647) at 37 °C for 5 min. To induce dimerization of the hook to the CD8, the medium was changed for DMEM with 200 nM rapamycin (Alfa Aesar) for 15 min at 37 °C. The cells were then fixed with fixation buffer (4 % formaldehyde, 4 % sucrose, 80 mM K-PIPES, 5 mM EGTA, 2 mM MgCl_2_, pH 6.8) for 10 min at RT. The coverslips were rinsed 4x 5 min with PBS and mounted in Mowiol and DAPI.

Cells were imaged using a spinning disc confocal system (Ultraview Vox; PerkinElmer) with a 100× 1.4 NA oil-immersion objective. Images were captured in Volocity using a dual-camera system (ORCA-R2; Hamamatsu) after excitation with lasers of wavelength 488 and 640 nm.

### Image analysis

The images acquired were duplicated and thresholded to isolate vesicular structures. To analyze only coinciding structures, thresholded images were multiplied with one another using the “Image calculator” plugin in FIJI and the vesicular structures measuring between 0.03 µm^2^ to 0.8 µm^2^ and of 0.3-1 circularity were counted in the resulting image using the “analyze particles” plugin. A one-way ANOVA with Dunnett’s post-hoc test was performed using GFP-FKBP as control.

### Buffer compositions

HKM buffer: 25 mM HEPES pH 7.2, 125 mM potassium acetate, 5 mM magnesium acetate. Tris buffer: 1 M Tris pH 7.1, 1 mM EDTA, 1 mM DTT. Ficoll/Sucrose buffer: 6.3 % w/v Ficoll PM 70, 6.3 % w/v sucrose in HKM pH 7.2. Saturated ammonium sulfate: excess ammonium sulfate dissolved in 10 mM Tris pH 7, 0.1 mM EDTA. Polymerization buffer: 100 mM MES pH 6.4, 1.5 mM MgCl_2_, 0.2 mM EGTA. Depolymerization buffer: 20 mM TEA pH 8.0, 1 mM EDTA, 1 mM DTT. Purification buffer: 20 mM HEPES pH 7.2, 200 mM NaCl. Elution buffer: 20 mM HEPES pH 7.0, 200 mM NaCl, 10 mM reduced glutathione. Prescission buffer: 50 mM tris-HCl pH 7.0, 150 mM NaCl, 1 mM EDTA, 1 mM DTT.

### Protein expression and purification

Clathrin was purified from pig brain clathrin-coated vesicles using previously described methods (Rothnie et al., 2011). Clathrin cages were assembled for harvesting by dialyzing the purified triskelia into polymerization buffer at 4 °C and then harvested by ultracentrifugation. Pellets containing clathrin cages were resuspended in a small volume of polymerization buffer. Concentration of clathrin cages was assayed by A_280_ of triskelia to avoid the effects from light scattering.

β2HA was expressed as a GST-β2HA fusion protein in *Escherichia coli* strain, BL21. Bateria were grown at 37 °C to an OD_600_ of 0.6 and then induced with 0.8 mM IPTG at 20 °C overnight. Cells were harvested and resuspended in purification buffer (supplemented with Complete Protease Inhibitor Cocktail tablet as per Roche Applied Science instructions), and lysed by sonication. The soluble fraction was obtained by centrifugation at 25 000 rpm for 30 min. GST-β2HA was purified from the soluble fraction using glutathione resin (GE Healthcare). The GST tag was removed by overnight cleavage at 4 °C using GST fusion 3C protease (Prescission, GE Healthcare). Fusion protease was removed using glutathione resin and the cleaved β2HA was collected in the flow-through. Cleaved β2HA was concentrated and loaded onto a HiLoad Superdex 200 (equilibrated in purification buffer) for further purification via size exclusion chromatography. Fractions containing purified β2HA were pooled and concentrated on Vivaspin columns (Sartorius).

### β2HA-clathrin complex preparation

To identify the maximum amount of β2HA that could bind the clathrin cages, increasing molar amounts of β2HA was reconstituted with 3 µM clathrin in depolymerization buffer at 4 °C and subsequent dialysis overnight into polymerization buffer at 4 °C. The β2HA-clathrin cage complexes were harvested by centrifugation at 230 000 *g* for 30 min and concentrated 10-fold by pellet resuspension into a small volume of polymerization buffer. The protein composition of the re-suspended pellets was analyzed by SDS-PAGE and densitometry in ImageJ (Schneider et al., 2012).

### Negative stain transmission electron microscopy

Clathrin cages reconstituted in the presence of 180 µM β2HA were screened under negative stain. Assembled β2HA-clathrin cage complexes (5 µl of 1 µM) were pipetted onto a glow-discharged formvar carbon 300-mesh copper grid (Agar Scientific) and incubated for 1 min at room temperature. Excess sample was removed by blotting with Whatman filter paper and 5 µl of 1 % (w/v) uranyl acetate stain was subsequently applied to the grid and left to incubate for 1 min at room temperature. Excess negative stain was removed by blotting with Whatman filter paper. Samples were imaged using a JEOL 2100Plus and Gatan OneView IS at 200 keV.

### Cryo-electron microscopy

3 µl of β2HA clathrin cage complexes (clathrin at 9 µM) were applied to glow-discharged 300-mesh copper Quantifoil R1.2/1.3 grids and blotted at 4 °C and >90 % humidity for 4.5 s before plunging into an ethane/propane mix (80 %/20 %) liquefied and cooled by liquid nitrogen using a Leica EM GP automated plunge freezing device.

Cryo-electron micrographs were collected as movies using a Titan Krios and Falcon III detector (Leicester Institute of Structural and Chemical Biology), operating at 300 keV. EPU was used for automated data collection, movies were acquired at a total dose of 64 e^−^Å^−2^ over 1 s at a dose rate of 1.65 e^−^ Å^−2^ s^−1^ with a magnified pixel size of 1.39 Å px^−1^ using a 1 µm beam and 70 µm C2 aperture. Three images were acquired per hole with some illumination of the carbon support. Micrographs were targeted for collection between 1.1 µm to 2 µm defocus.

### Data processing

Beam-induced motion of the specimen was anisotropically corrected, with and without dose-weighting, using MotionCor2 (Li et al., 2013). The contrast transfer function of the motion-corrected summed micrographs was estimated from non-dose weighted micrographs using gctf v1.06 (Zhang, 2016) employing equiphase averaging and validation functions. RELION v3.0.5 (Scheres, 2012) was used for particle picking, extraction, and all classifications and refinements. 57 528 particles were manually picked from the non-dose weighted, motion-corrected micrographs and then extracted at a binned pixel size of 10.8 Å px^−1^. Reference-free 2D classification, over 25 iterations, was first used to analyze the quality of the extracted particles. The highest quality classes, containing 51 133 particles, were selected for further 3D classification. As previously described, supervised asymmetric 3D classification successfully sorted the particles into ten structural classes (Morris et al., 2019). The particles associated with the minicoat cage type, which produced the highest quality 3D classification output, were selected for subsequent hierarchical, supervised 3D classifications to identify the particles most stably associated with this particular cage geometry (Supplementary Figure S2). Further unsupervised 3D classification of these stable minicoat particles subdivided the particles into 3 classes and was carried out using a regularization parameter (*T*) of 4, no imposed symmetry and no mask (Supplementary Figure S3). 3D auto refinement of the most stable minicoat particles (at 10.8 Å px^−1^, without symmetry imposed) yielded a 24 Å minicoat volume. These particles were re-extracted from non-dose-weighted micrographs with a box size of 500 px and a pixel size of 2.78 Å px^−1^; large enough to include clathrin cages over 1000 Å diameter. The minicoat cage architecture was refined at 2.78 Å px^−1^ (i.e. binned two-fold) without imposing symmetry. The refinement reference (from the previous 3D auto refinement of minicoat particles) was low pass filtered to 40 Å. Since the output volume was a minicoat with mixed handedness, an unsupervised 3D classification was conducted on the minicoat particles (no symmetry imposed, and no alignment of particles). Only the minicoat particles producing contributing to volumes that had 100 % surface density were saved and used in subsequent processing. These particles were refined as described for the previous 3D auto refinement, and yielded a 11.7 Å volume. A mask was generated from this C_1_ reconstruction at 3σ, extended and softened by 2 and 9 px. This mask was employed in subsequent C_1_ refinements that used dose-weighted minicoat particles, solvent flattening and a Gaussian noise background. Reconstructions with and without imposed symmetry correlated well.

Resolution of each reconstruction was estimated using the gold-standard Fourier shell correlation (FSC) measurement within a mask created from the refinement volume (using threshold value of 3σ, expanded by 2 px to 4 px and softened by 9 px). The MTF of the Falcon III camera (operated at 300 keV) was applied and the B factor of the map was automatically calculated if the resolution exceeded 10 Å. In instances, where sub-10 Å refinements were calculated, a user-defined B factor value was given.

In order to identify β2HA in the minicoat volume we subtracted the signal contributed by the outer coat region and subsequently conducted a masked, unsupervised 3D classification on the signal-subtracted inner cage region (Supplementary Figure S5) with a regularization parameter (*T*) of 20 tetrahedral (T) symmetry imposed and no alignment of particles calculated. The mask was created from the tetrahedral refinement volume using a threshold value of 3σ, expanded by 5 px and softened by 9 px. The particles contributing to each of the 20 classes were saved separately and refined with *T* symmetry imposed. Qualitative analysis of the individual refinement outputs (visualized at contour level 3σ), identified two classes that possessed strong additional density that was not present in reconstructions calculated using signal from the whole cage (i.e. prior to signal subtraction) *or* in a minicoat cage volume reconstructed *without* adaptor protein present.

### Localized subparticle extraction and reconstruction

To improve the resolution of the strong additional density resolved after masked 3D classifications of signal-subtracted minicoat particles, we performed localized reconstruction (Ilca et al., 2015) as previously described for single particle data sets of clathrin cages (Morris et al., 2019). Hub regions were extracted and recentered as new subparticles in 350 px boxes from whole minicoat cage particles. Each of the extracted hub subparticles were reconstructed separately to serve as references in subsequent refinements.

All refinements were conducted in C_1_ with masking applied from a 3σ extended 2 px and softened 9 px mask (3σ/e2/s9). Global resolution of the hub region was estimated as described previously using the gold-standard FSC approach (within a mask 3σ/e2/s9). The refinement was found to have converged at 9.6 Å. Local resolution estimations were made using ResMap (Kucukelbir et al., 2014) revealing lower resolutions in the terminal domain regions of the minicoat hub. To improve the quality of the β2HA density located between the terminal domains under the hub vertex, the hub subparticles were classified based on whether the β2HA density connected terminal domains from two separate pentagonal faces (PP) or connected a hexagonal and pentagonal face (HP). Compared to the whole cage volume (post-signal subtraction), the resolution of the β2HA (and neighboring clathrin heavy chain regions) is improved, allowing PDBs of the clathrin heavy chain (residues 1-361, 362-487 and 488-834) to be fitted into the hub volume.

### Global difference analysis

Student’s t-test was used to determine the significance of differences between two structures, using SPIDER and the programs of Milligan and Flicker as previously published (Frank et al., 1996; Milligan and Flicker, 1987; Young et al., 2013). In order to do this, independent maps of each structure (four in the case of whole cages, three for the hubs) were created using Relion, by dividing the data (by splitting the star files using `-split` command in `relion_star_handler` script (in RELION) into separate sets taking care to distribute images of particles evenly from each micrograph and therefore the defocus spread. A low pass Fourier filter, 11 Å in the case of whole cages and 12 Å for the hubs, was applied to the maps. In order to avoid potential false differences due to differences in the quality of the two structures solved, or random effects such as the sampling of defocus values, the two structures were scaled together in reciprocal space by calculating their radial amplitude-profiles. A reciprocal-space scaling profile was calculated by comparing the amplitude profile of the clathrin-only map with the β-adaptin map (Young et al., 2013). Using this, all the β-adaptin sub-maps were rescaled to fit the profile of the clathrin-only map. These maps were used to calculate an average and variance for each structure. The per voxel value of *t* and the significance of differences was computed from these, using the appropriate degrees of freedom. Many regions had significant differences with *p <* 0.05. Regions we have interpreted to show direct density differences relating to ligand binding have *p <* 0.0001. The images show the original maps, with the value of *p* colored onto the surface according to the scale shown.

### Redocking the adaptor proteins

Initially, the β2-appendage protein structure (1E42) was docked by fitting into the unoccupied density in the β2HA-clathrin map. However, this led to an overlap of residues between the C-terminal domain and its neighboring terminal domain, while leaving a gap at the putative interface between the N-terminal domain and a second local terminal domain. The docking program BUDE (McIntosh-Smith et al., 2012, 2015) was used to refine this structure as follows. The complex was centered on the center-of-coordinates of the β2-appendage and the complex split into clathrin as the receptor and β2-appendage as the ligand. The docking grid was defined: −10 to 10 in 2° increments for rotation and −10 to 10 in 1 Å increments for translation. A genetic algorithm, EMC (Abraham et al., 2015), sampling 1.1 million poses was used to find low energy poses. The best pose was inspected and new rotamers chosen for a few interfacial sidechains to optimise putative interfacial interactions (beta-ear: R732, R759, N846, E849, E882; clathrin: R188) and the above docking procedure repeated. Next, Gromacs 2019.4 (Abraham et al., 2015) was used to parameterise the complex with the Amber99SB-ildn (Lindorff-Larsen et al., 2010) forcefield at pH 7 and place it in box of TIP3P water containing 0.15 M NaCl. A short energy minimization (200 steps of steepest descents) was performed to remove bad intermolecular atom-atom contacts, permitted by BUDE’s very soft empirical free energy forcefield, and yield the finished model.

The initial α-appendage complex was prepared by superimposing the α-appendage (1B9K) on the β2-appendage C-terminal domain of the finished complex. This preliminary model of the α-appendage complex was subjected to the same docking and minimization procedure described above. Because the angle between the platform and sandwich domains of the α-appendage is smaller, we also prepared complexes where either the platform or sandwich domain of the α-appendage (1B9K) were directly superimposed onto the corresponding region of the β2-appendage domain of the finished complex using the Matchmaker function in UCSF Chimera.

### *in silico* alanine scanning

The two energy-minimized complexes and the individual platform and sandwich domain α-appendage complexes were presented to the BAlaS server http://balas.app (Wood et al., 2020). The three clathrin domains were assigned as the receptor and the appendages (either β2 or α) as the ligand. Alanine scanning and constellation calculations were performed and the results downloaded.

## Figure preparation

Maps and models were visualized using UCSF Chimera (Pettersen et al., 2004). Simplified views of triskelia were generated using IgorPDB and 3IYV. Microscopy figures and plots were made Fiji and Igor Pro (WaveMetrics Inc.). All figures were assembled in Adobe Illustrator.

## Data Availability

EM maps supporting this study have been deposited in the Electron Microscopy Data Base with accession numbers 12980.emd, 12981.emd, 12983.emd and 12984.emd (relating to Figures 3, 4 B and C, 4 A and 5, respectively). The fitted model of clathrin terminal domains and β2-appendage has been deposited in the protein data bank as 7OM8.pdb. Original models of both the clathrin terminal domain [1BPO.pdb, (ter Haar et al., 1998)] and β2-appendage [1E42.pdb, (Owen et al., 2000)] were used to generate the fitted model with only the interfaces between the protein molecules re-modelled. No inter-molecule clashes have been identified in the fitted model. The intra-molecule clashes and geometric outliers are historical

## ACKNOWLEDGEMENTS

SMS and CJS thank BBSRC for support (grant BB/N008391/1). CJS was a Royal Society Leverhulme Trust Senior Research Fellow. We acknowledge the Midlands Regional Cryo-EM Facility at the Leicester Institute of Structural and Chemical Biology (LISCB), major funding from MRC (MC_PC_17136) and thank Christos Savva and T. J. Tagan for assistance with data collection. Sample preparation and development was supported by Saskia Bakker, Warwick Advanced Bioimaging Research Technology Platform, using equipment funded by BBSRC ALERT14 award BB/M01228X/1 and MRC award reference MC_PC_17136. We thank Laura Wood and Miguel Hernández González for early work on this project.

## AUTHOR CONTRIBUTIONS

SMS carried out structural biology experiments and contributed to the manuscript writing and figure preparation. GL conducted hot wiring experiments contributed to the manuscript writing and figure preparation. KMW carried out structural biology experiments and contributed to the manuscript writing. KLM contributed to structural analysis and manuscript writing. AMR carried out structure comparisons and contributed to the manuscript writing. RBS carried out model building and contributed to the manuscript writing. SJR contributed to data interpretation, manuscript writing and figure preparation. CJS contributed to data interpretation, structural analysis and wrote the final draft, which was approved by all authors.

## Supplementary Information

**Figure S1.**
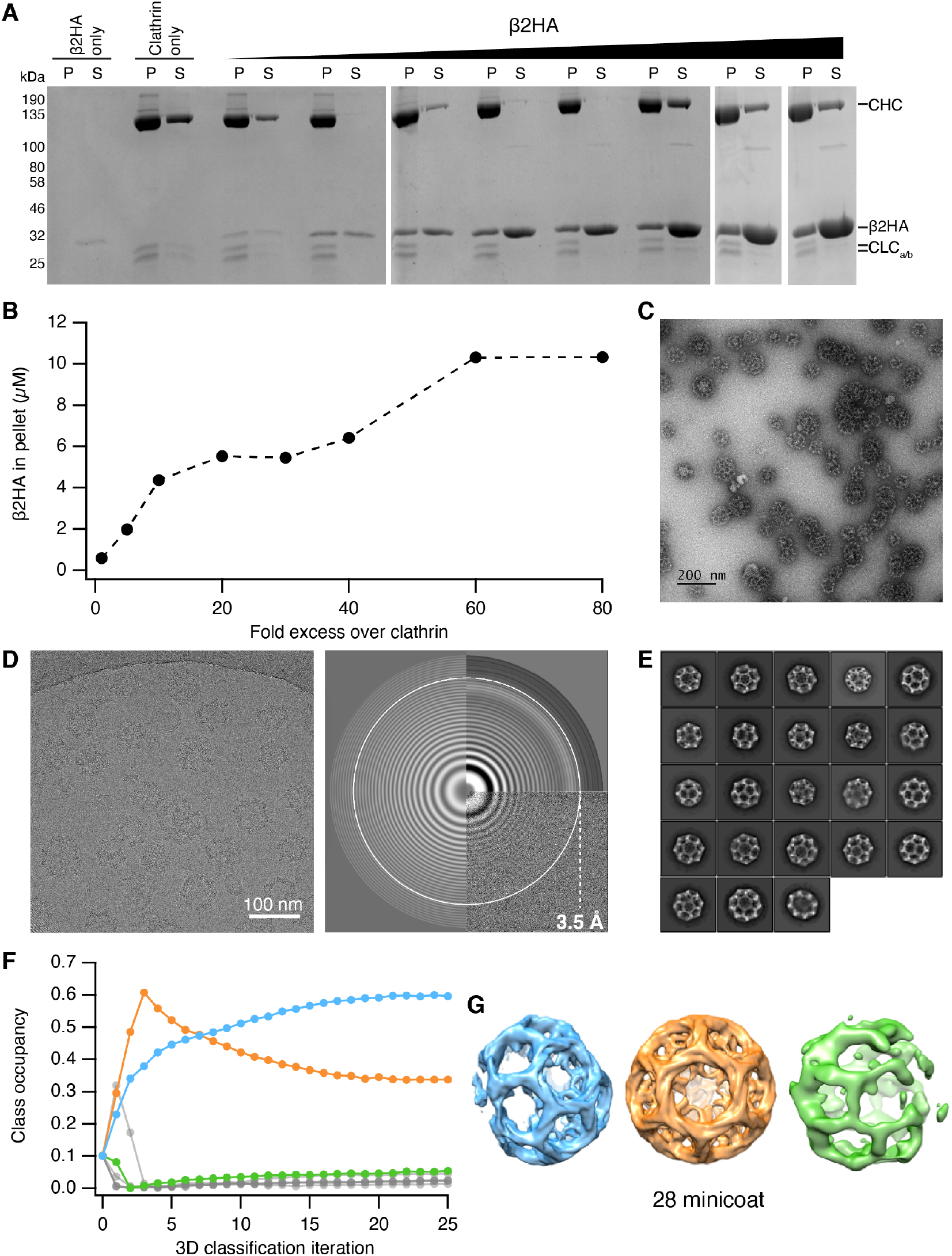
Clathrin-β2HA reconstitution, data collection and processing. (**A**) Clathrin triskelia (3 µM) were assembled in the presence of increasing concentrations (3 µM to 240 µM) of β2-adaptin_616-951_(β2HA). Clathrin assemblies were pelleted and analysed by SDS-PAGE to determine the amount of clathrin (CHC and CLCa/b) and β2HA in the pellet (P). (**B**) Densitometry of gels in A shows that increasing amounts of β2HA pelleted with clathrin during the reconstitution experiments, with a 60-fold excess of adaptor protein yielding the maximum amount of clathrin-binding. (**C**) Negative stain TEM analysis of clathrin cages reconstituted with a 60-fold excess (240 µM) of β2HA. Scale bar = 200 nm. (**D**) A representative cryo-electron micrograph (left) at −1.4 µm defocus and the corresponding power spectrum indicating the information content at high spatial frequencies (right). (**E**) 2D class averages of classes that were selected for 3D classification in RELION. (**F**) Particle occupancy of the 10 classes obtained with supervised, asymmetric 3D classification in RELION. (**G**) 3D surface representations of the 3 clathrin cages generated from the supervised, asymmetric 3D classification of clathrin cage particles. Their color corresponds to the class occupancy data shown in panel F. Only the orange cage was reconstructed in full, enabling its cage geometry to be confirmed as minicoat.

**Figure S2.**
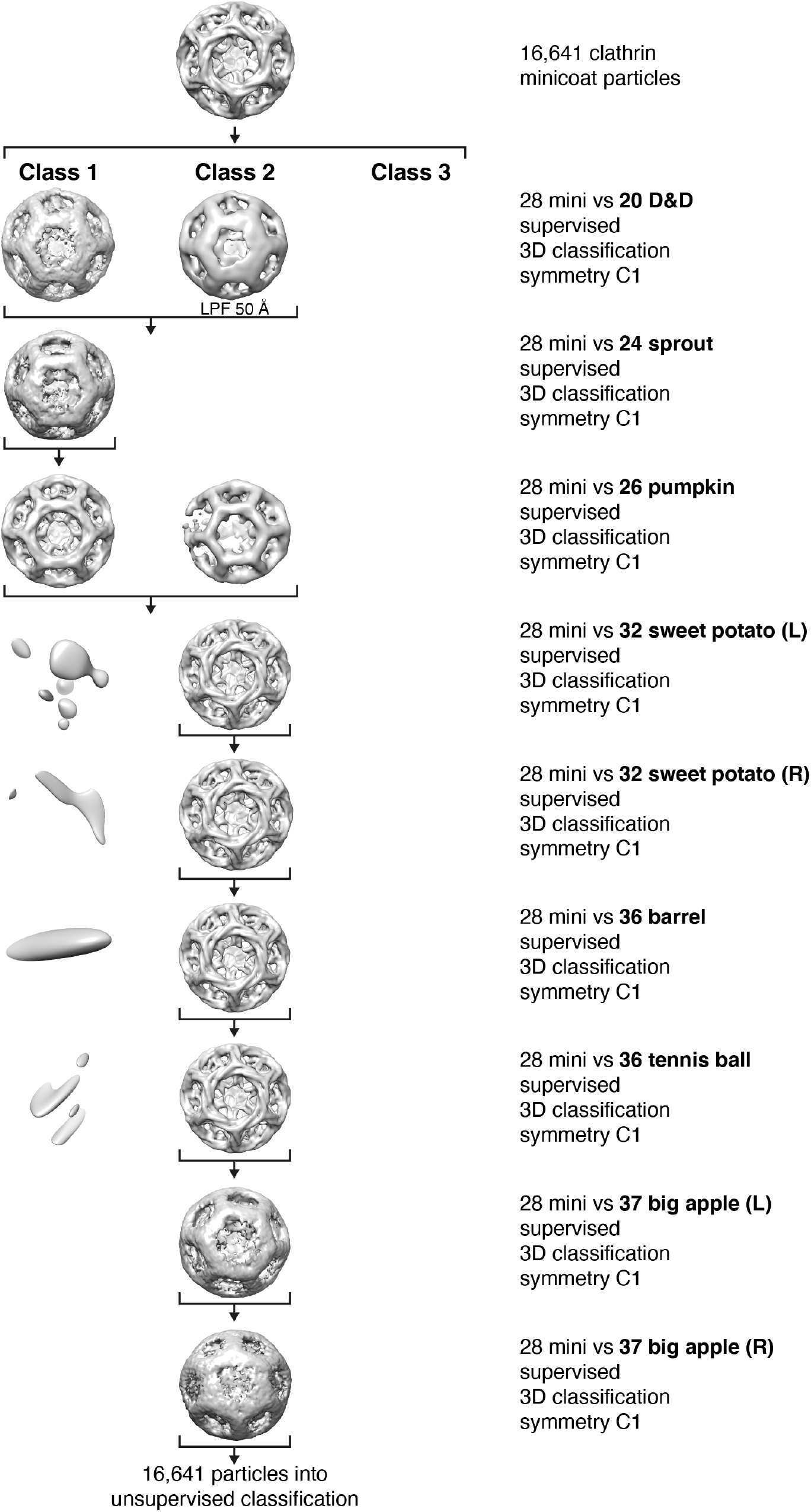
Supervised, hierarchical 3D classification scheme in RELION. To identify particles most stably associated with the minicoat cage type, the 16 641 clathrin cage particles which were classified as a minicoat cage type during a supervised, asymmetric 3D classification (see Supplementary Figure S1), were input into a hierarchical 3D classification scheme in RELION. A total of 9, separate, 3D classifications were conducted whereby particles were classified against 2 reference structures: a minicoat and one of the nine other geometric references in the cage library. Only the particles classifying as a minicoat cage type were retained and input into the next 3D classification (indicated by arrows). If the 3D output was of poor quality, the volume was low-pass filtered (LPF) to aid identification of cage architecture. The output of each classification can be seen in each row of the figure.

**Figure S3.**
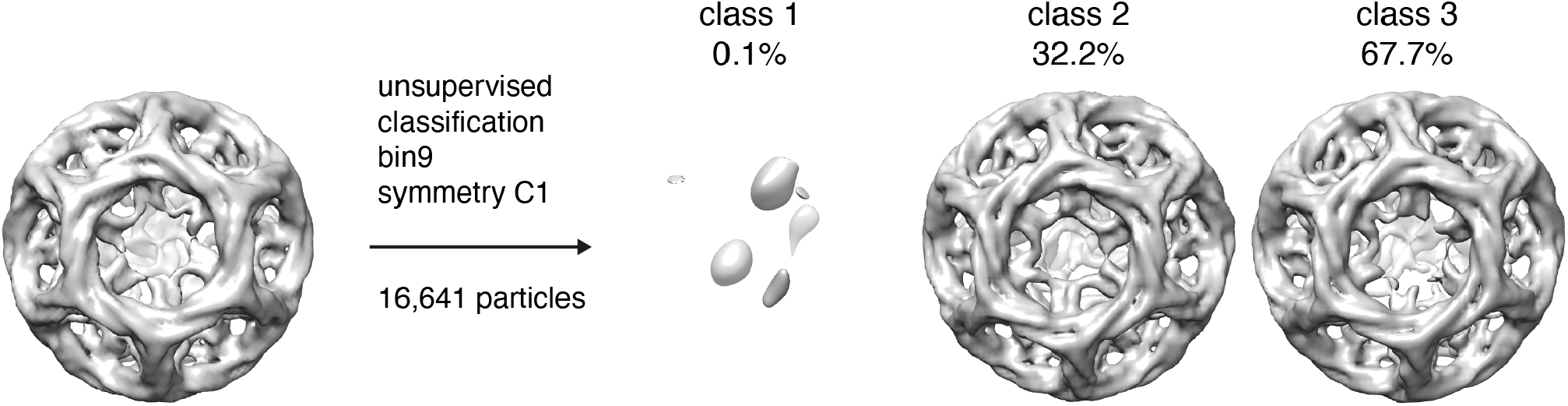
Unsupervised 3D classification in RELION. Unsupervised classification of 16 641 minicoat particles. Output reconstructions were compared with the input supervised 3D classification volume. Aside from opposite handedness, classes 2 and 3 were architecturally identical to the input volume.

**Figure S4.**
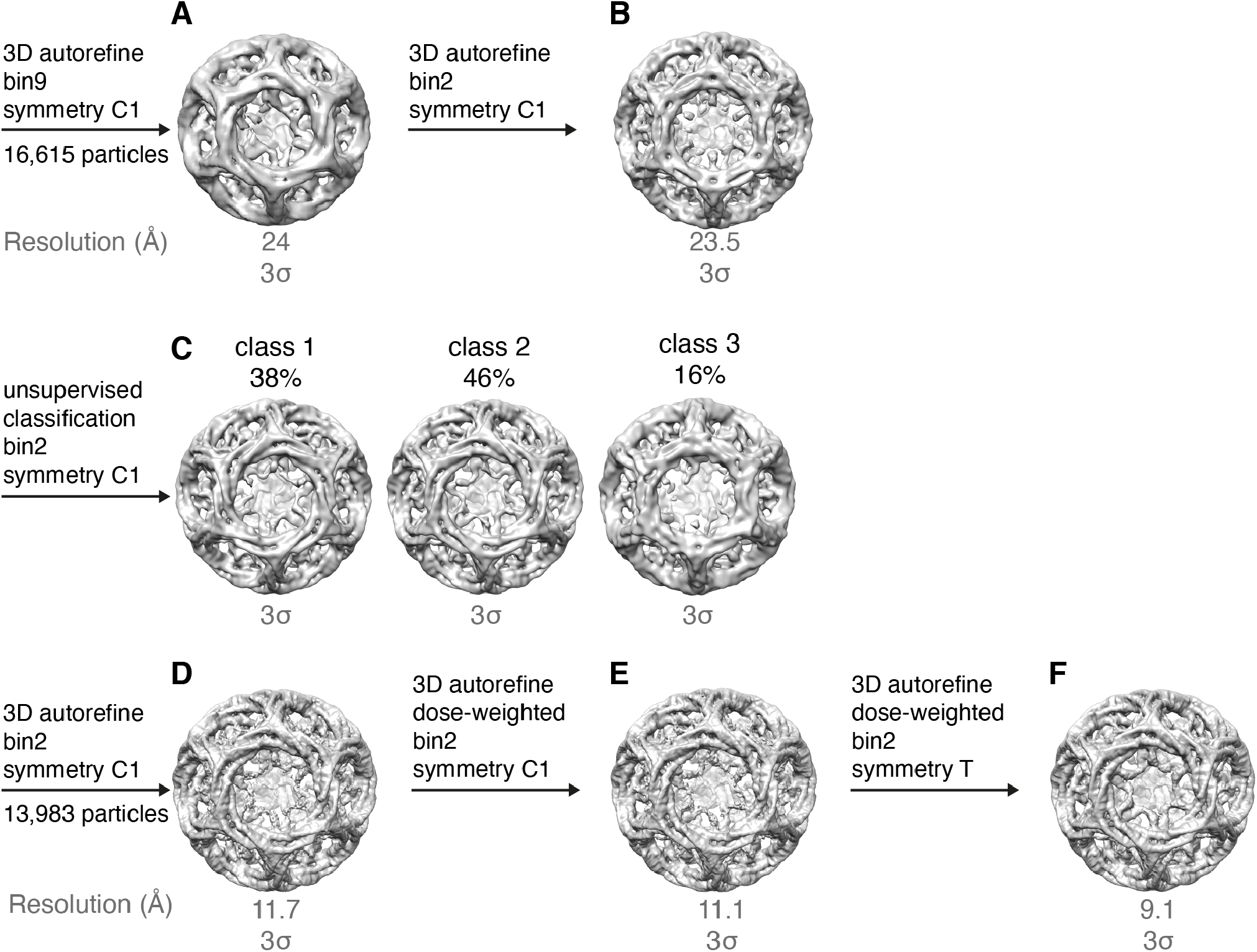
Image processing of minicoat cage particles in RELION. (**A**) Initial 3D auto refinement of 16 615 minicoat particles yielded a 24 Å volume with minicoat architecture. (**B**) Unbinning of the particles from A to 2.78 Å px^−1^ and input into another 3D auto refinement produced a minicoat volume with mixed handedness. (**C**) To identify any minicoat particles of poor quality, particles from B were input into an unsupervised 3D classification. Class 3 contained particles that failed to produce a whole minicoat cage structure; therefore, only particles from classes 1 and 2 were taken to the next stage of image processing. D, E and F represent the final 3 rounds of 3D auto refinement with input parameters stated on the figure. A 9 Å minicoat cage structure was the final output.

**Figure S5.**
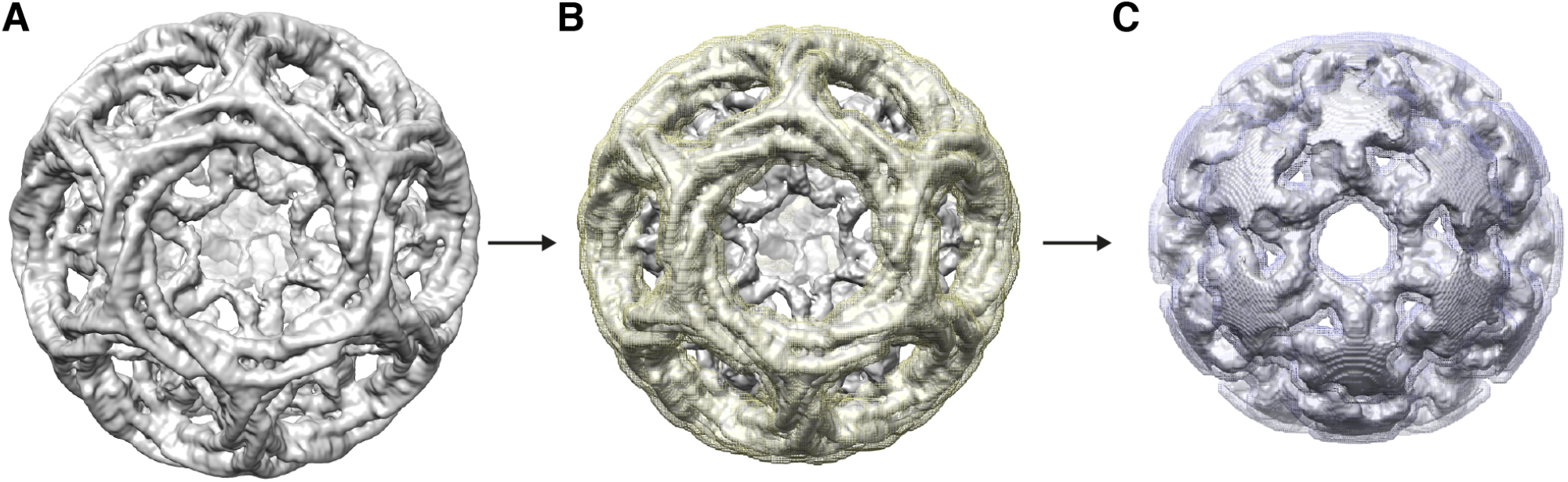
Signal subtraction of outer minicoat cage region. (**A**) 9 Å minicoat cage structure at 3σ contour level. A soft mask (marked yellow in B) was applied to A to identify the protein density to be subtracted. The remaining density was input into a masked (marked purple in C), unsupervised 3D classification in RELION.

**Figure S6.**
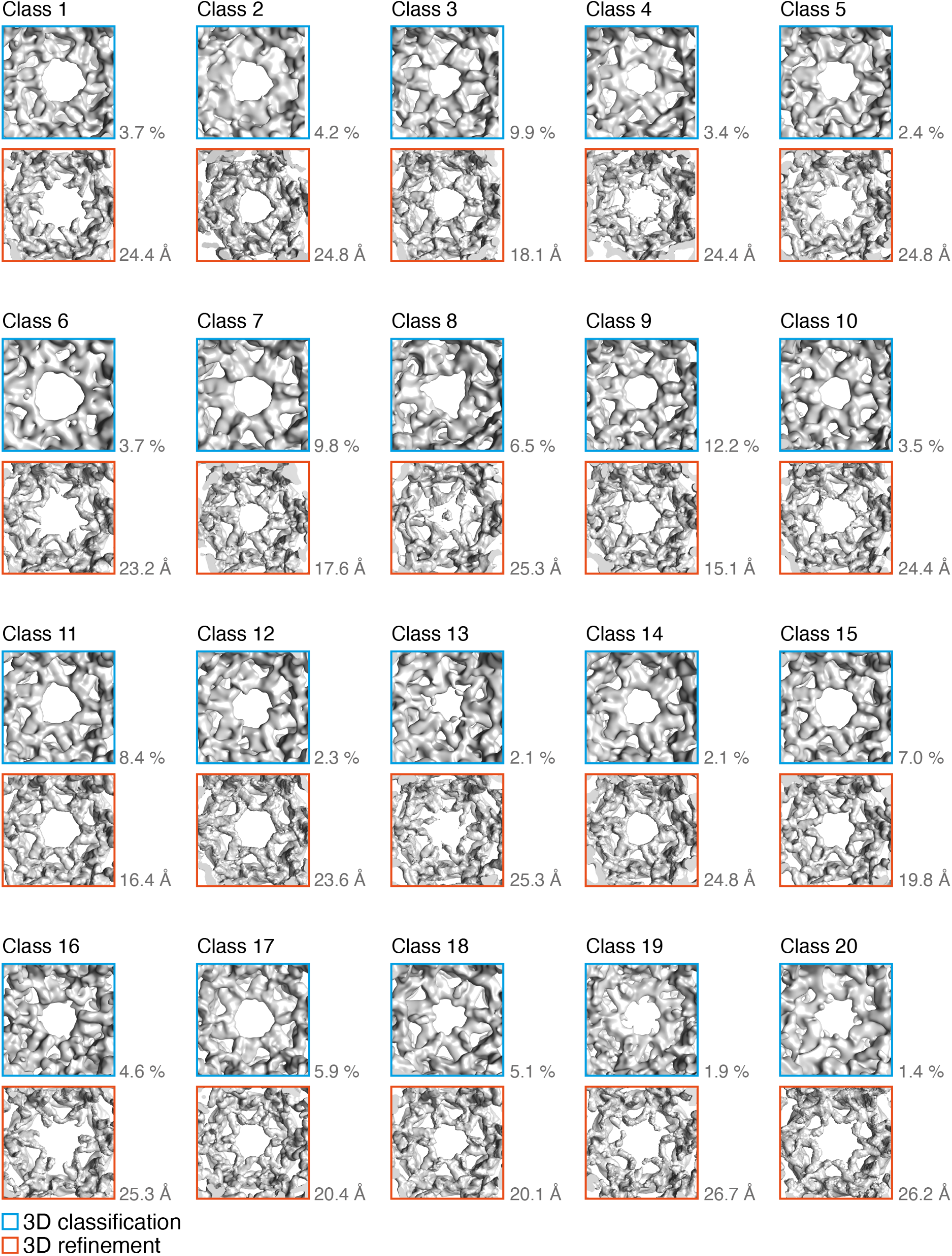
Unsupervised, masked 3D classification of signal subtracted minicoat cage particles. Output of unsupervised, masked 3D classification of signal-subtracted minicoat cage particles. Particles were separated into 20 classes: hexagonal faces (representative of average class quality for all remaining polygonal faces) are shown for each class with the corresponding refinement (at 3σ contour level) shown below.

**Figure S7.**
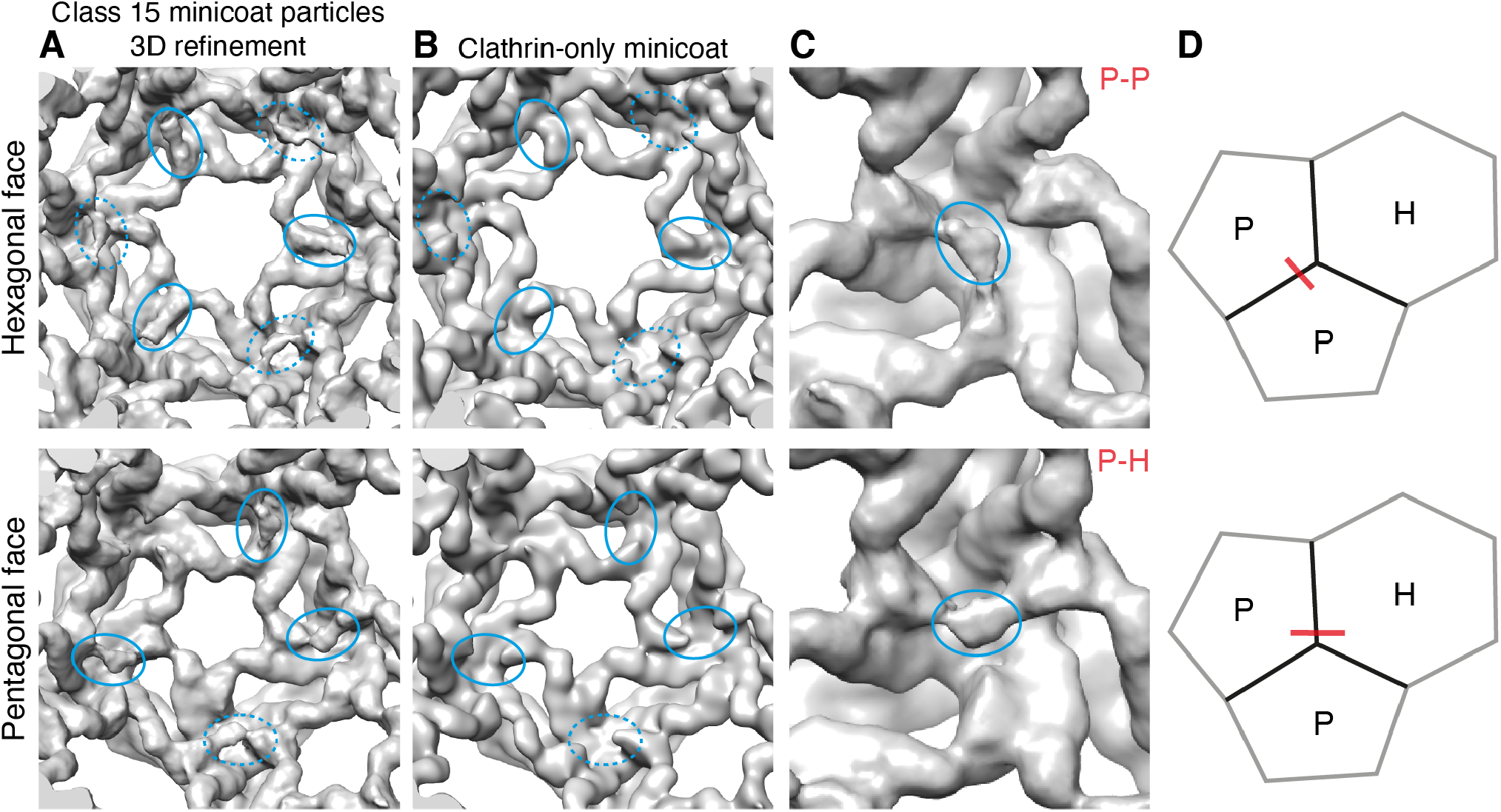
Locating and sub-classifying β2HA density in class 15 of masked, 3D classification output. (**A**) Representative hexagonal and pentagonal faces for class 15 3D auto refinement. Solid blue ellipses highlight new density seen following masked, 3D classification within a given polygonal face. Dashed ellipses highlight densities connecting adjacent polygonal faces. (**B**) Representative hexagonal and pentagonal faces for clathrin-only minicoat cage (low-pass filtered to 20 Å). Equivalent positions to those in column A are marked in ellipses, highlighting the lack of density in these regions. (**C**) density cross-linking terminal domains from two, adjacent pentagonal faces (denoted P-P) is marked in blue ellipse. Density cross-linking terminal domains from adjacent hexagonal and pentagonal faces (denoted P-H) is marked in blue ellipse. The geometric context of P-P and P-H densities is depicted in **D**.

**Figure S8.**
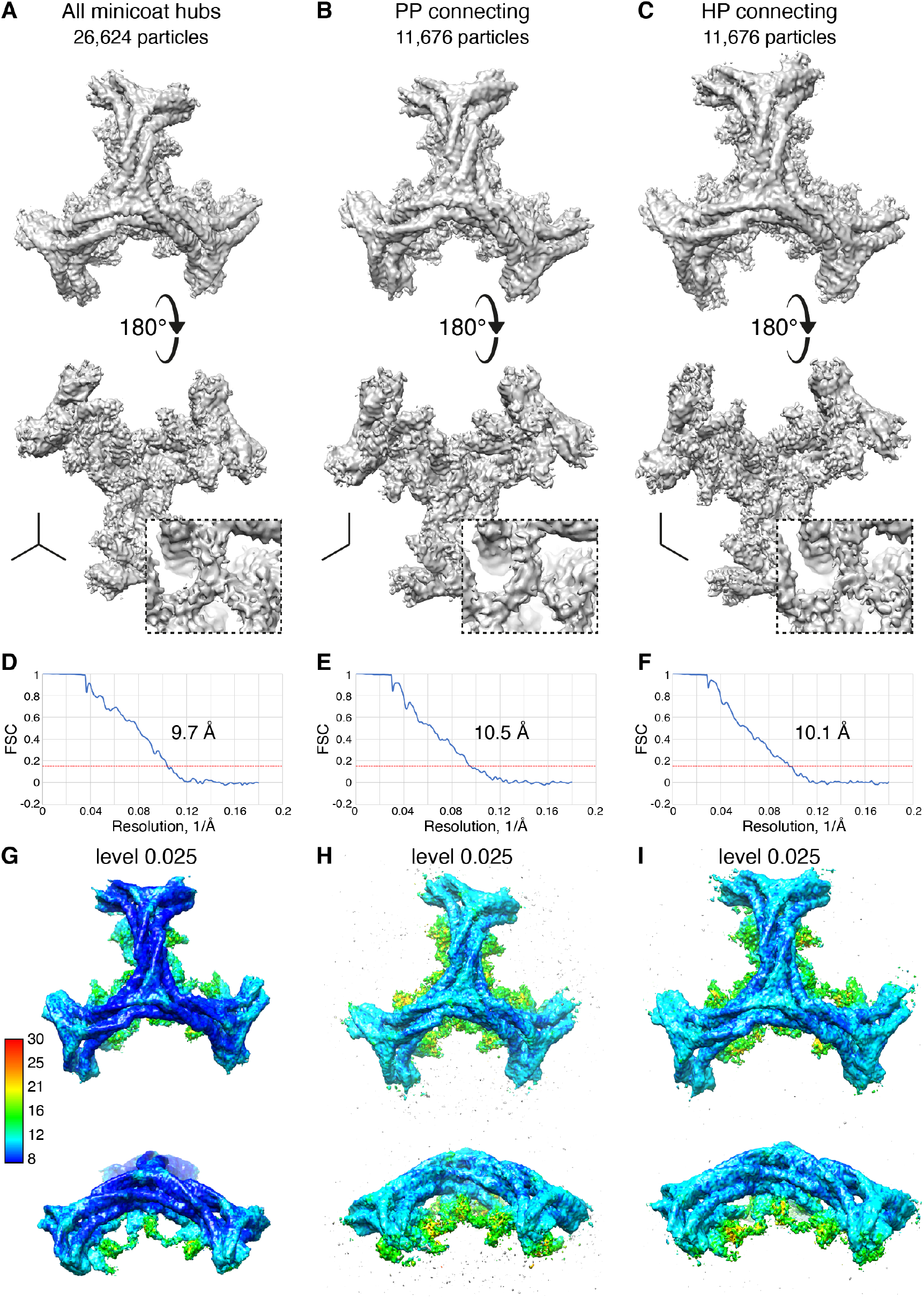
Localized reconstruction of minicoat cage particles. (**A**) Asymmetric 3D auto refinement of hub regions for all 26 624 minicoat particles from class 15 yielded a 9.7 Å resolution volume. 180°rotation of this asymmetric unit revealed a single β2-appendage connecting two or three terminal domains (panel below and inset). Local resolution of this region ranged from 12 Å to 16 Å (G). (**B** and **C**) Sub-classification of hub particles based on geometric context. P-P and H-P hubs (defined in Supplementary Figure S7) yielded volumes resolved at 10.5 Å and 10.1 Å volumes, respectively. The local resolution of the lower hub regions was approximately 16 Å (H and I). Both volumes gave improved definition of the β2-appendage density. (**D, E** and **F**) FSC plots for maps shown in A, B and C, respectively. The resolution cut-off used was at a correlation value of 0.143. (**G, H** and **I**) Shows maps A, B and C respectively coloured by local resolution. The scale gives the local resolution in Å.

**Figure S9.**
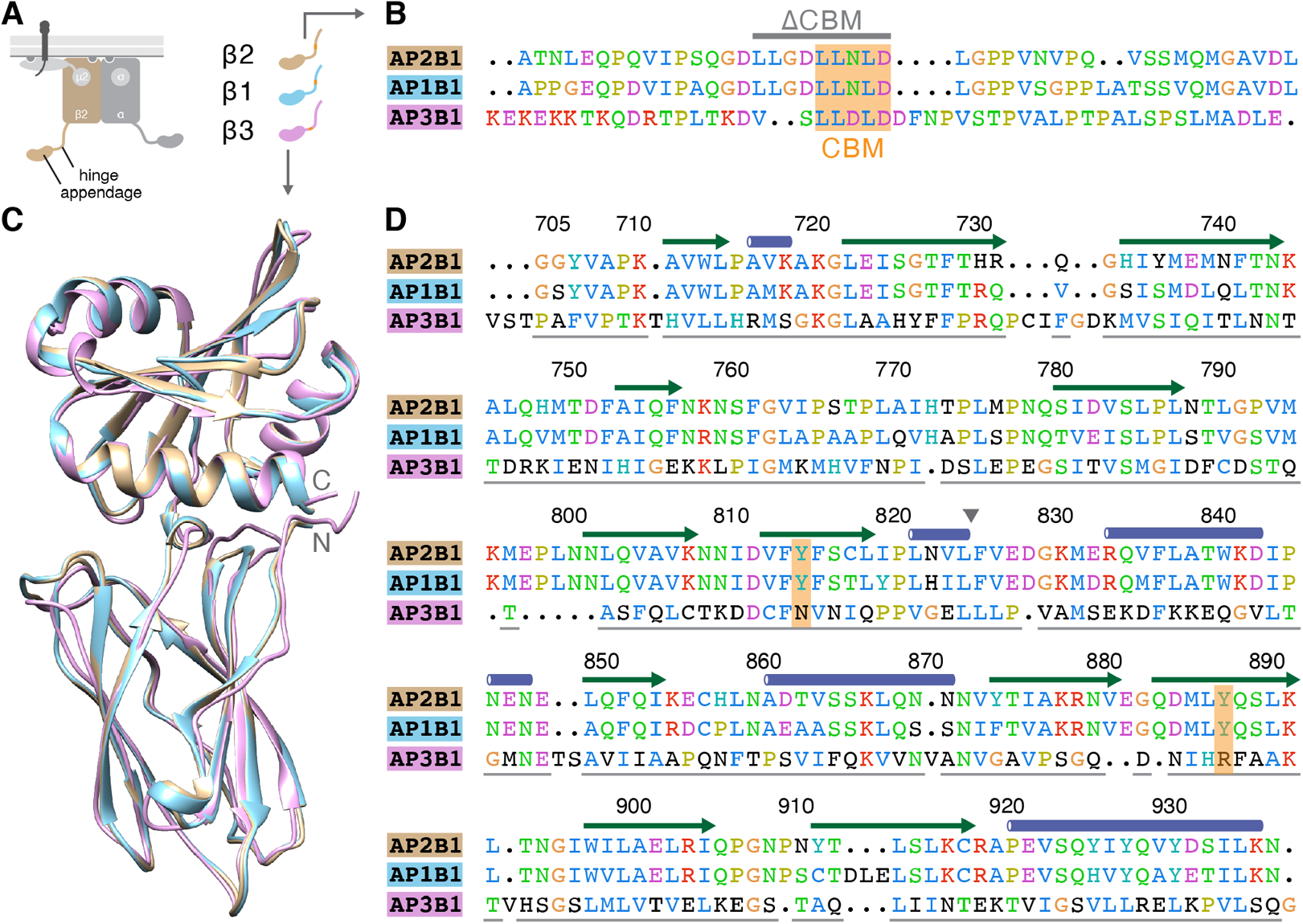
Sequence comparison between β2 and β1 or β3. (**A**) Schematic diagram of the AP2 complex showing the position of hinge and appendage of the β2 subunit. The color coding for hinge and appendage of β2, β1 and β3 is shown (left) with the position of the clathrin-box motif (orange). (**B**) Alignment of a section of the hinge region of β2, β1 and β3 containing the LL[D/N]LD clathrin-box motif (orange, CBM). The region deleted in ΔCBM construct is indicated by a gray line. Residues, β2 612-655, β1 613-658 and β3 835-883, are colored according to property. (**C**) Overlay of β2, β1 and β3 appendage structures. The β2 structure is PDB code 2G30, β1 was created using MODELLER with 2G30 as a template and β3 was created using I-TASSER using 1E42 as a template. (**D**) Alignment of the appendage regions shown in C. Structural features and numbering of β2 is shown above. Triangle indicates the separation between the two lobes of the appendage. The position of Tyr 815 and Tyr 888 is indicated in orange. Gray lines indicate structural alignment. Residues are shown colored by property, black residues are shown against consensus.

